# Potent and lasting seizure suppression by systemic delivery of antagomirs targeting miR-134 timed with blood-brain barrier disruption

**DOI:** 10.1101/797621

**Authors:** C. R. Reschke, L. F. A. Silva, V. R. Vangoor, M. Rosso, B. David, B. L. Cavanagh, N. M. C. Connolly, G. P. Brennan, A. Sanz-Rodriguez, C. Mooney, A. Batool, C. Greene, M. Brennan, R. M. Conroy, T. Rüber, J. H. M. Prehn, M. Campbell, R. J. Pasterkamp, D. C. Henshall

## Abstract

RNA therapies such as oligonucleotides (OGNs) offer precision treatments for a variety of neurological diseases, including epilepsy but their deployment is hampered by the blood brain barrier (BBB). Here we used brain imaging and assays of serum proteins and tracer extravasation, to determine that BBB disruption occurring after status epilepticus in mice was sufficient to permit passage of systemically-injected antisense OGNs targeting microRNA-134 (Ant-134) into the brain parenchyma. A single intraperitoneal injection of Ant-134 two hours after status epilepticus in mice resulted in potent suppression of spontaneous recurrent seizures, reaching a 99.5% reduction during recordings at three months. The duration of spontaneous seizures, when they occurred, was also reduced in Ant-134-treated mice. These studies indicate that systemic delivery of Ant-134 reaches the brain and produces disease-modifying effects after systemic injection in mice when timed with BBB disruption and may be a clinically-viable approach for this and other disease-modifying microRNA therapies.

## Introduction

Targeting coding and noncoding RNAs offers unprecedented potential for precision therapeutics and disease-modification (*1*). Among leading strategies is the use of antisense oligonucleotides (OGNs) which are now approved for several neuromuscular diseases and in clinical trials for other conditions (*1, 2*). There has been less progress, however, with deployment of OGNs for the treatment of brain diseases. This is in part due to the blood brain barrier (BBB) which prevents the passage of large macromolecules and small negatively charged molecules from the systemic circulation into the brain (*3*). Overcoming the BBB may be possible using encapsulation techniques or conjugation to cell-penetrant peptides, however the safety and efficacy of these approaches remain uncertain.

Epilepsy is a common brain disease characterised by recurrent spontaneous seizures. Many patients do not achieve seizure freedom with currently available anti-epileptic drugs and there is an urgent and unmet need for disease-modifying or anti-epileptogenic therapies (*4, 5*). MicroRNAs (miRNAs) have recently emerged as a novel class of therapeutic target in epilepsy (*6*). They are small noncoding RNAs that function post-transcriptionally to reduce protein levels via base-pairing to complementary regions in target mRNAs. In the brain, miRNAs are required for cell growth, differentiation and synaptic plasticity as well as the control of neuroinflammation and apoptosis (*7*). Notably, many of these processes are dysregulated in epileptogenesis (*6*). Altered miRNA biogenesis has been detected in the hippocampus of patients with epilepsy (*8*) and experimental deletion of certain miRNAs leads to neurodegeneration and epilepsy in mice (*9, 10*).

Increased brain levels of miR-134-5p (*MIR134*; hereafter, miR-134) have been widely reported in experimental and human temporal lobe epilepsy (*11-13*). miR-134 is a conserved, brain-enriched miRNA that controls dendritic spine morphology via regulation of LIM kinase-1 (Limk1) (*14*), a well-characterized serine/threonine kinase that can regulate actin polymerization, although other targets are known (*15, 16*). Antagomirs are modified antisense OGNs which produce potent and specific inhibition of miRNAs *in vivo* (*1, 2, 17*). Previous studies showed that reducing levels of miR-134 in the brain protects against evoked seizures in multiple chemoconvulsant models (*11-13, 18*). Moreover, intracerebroventricular (ICV) injection of miR-134-targeting antagomirs after status epilepticus produced potent and lasting reductions in the later occurrence of spontaneous recurrent seizures (SRS) in rats and mice (*11-13*). The mechanism of action is uncertain although silencing miR-134 *in vivo* produces dendritic spine phenotypes and rescue of protein levels consistent with a mechanism involving de-repression of Limk1 (*11, 18*).

As with other OGNs, studies tracking radiolabelled antagomirs have demonstrated they do not reach the brain after systemic injection in healthy animals (*19*). However, antagomirs targeting miR-134 (Ant-134) might be used therapeutically after acute brain injuries or status epilepticus. In this setting, there is likely to be a temporary disruption of BBB integrity (*20-22*). We hypothesized that we could deliver Ant-134 to a brain target (hippocampus) after systemic administration if injected concurrent with BBB disruption. Here we tracked BBB disruption in a mouse model of status epilepticus to inform the timing of systemic delivery of Ant-134. We followed the effects on epilepsy and show potent and lasting suppression of seizure activity and link the mechanism to de-repression of Limk1.

## Results

### Timing and size selectivity of BBB disruption following status epilepticus in mice

We first established the timing of BBB disruption after experimental status epilepticus and if it is sufficient to allow passage of Ant-134 into the brain after systemic injection. Status epilepticus was triggered by intraamygdala microinjection of kainic acid (KA) in mice, a well-characterised model of prolonged seizures that involves limbic networks and produces damage to the ipsilateral hippocampus followed by the emergence of SRS within 3 - 5 days (*23*). The model has proved apt to study anti-epileptogenic therapies and was previously used to study the effects of ICV-delivered Ant-134 on SRS in mice (*11*). Although studies have suggested the model is associated with BBB disruption (*24*), the timing and size selectivity of this disruption remains unknown.

We first confirmed macroscopic leakage of the BBB by intravenous (IV) injection of mice with Evans Blue, an azo dye that has a high affinity for serum albumin, and visualising extravasation at different time points (*20*). Whole and sectioned brains from these mice displayed Evans Blue staining in multiple brain regions after status epilepticus including the hippocampus (Fig. 1A). To support this finding, we used immunoblotting to assess the extravasation of serum proteins into brain tissue. Hippocampal samples from saline-perfused naïve mice or mice that received an intraamygdala injection of vehicle were largely devoid of the abundant serum proteins albumin and IgG (Fig. 1B-D; fig. S1A-B). In contrast, both proteins were readily detected in hippocampal lysates obtained as early as 1 hour after status epilepticus, consistent with BBB disruption (Fig. 1B-C; fig. S1A-B). A peak of extravasation was observed at 2 hours after status epilepticus (Fig. 1B-D), and both albumin and IgG remained detectable in hippocampal lysates at later time points (Fig. 1B-C; fig. S1B).

**Figure 1.**
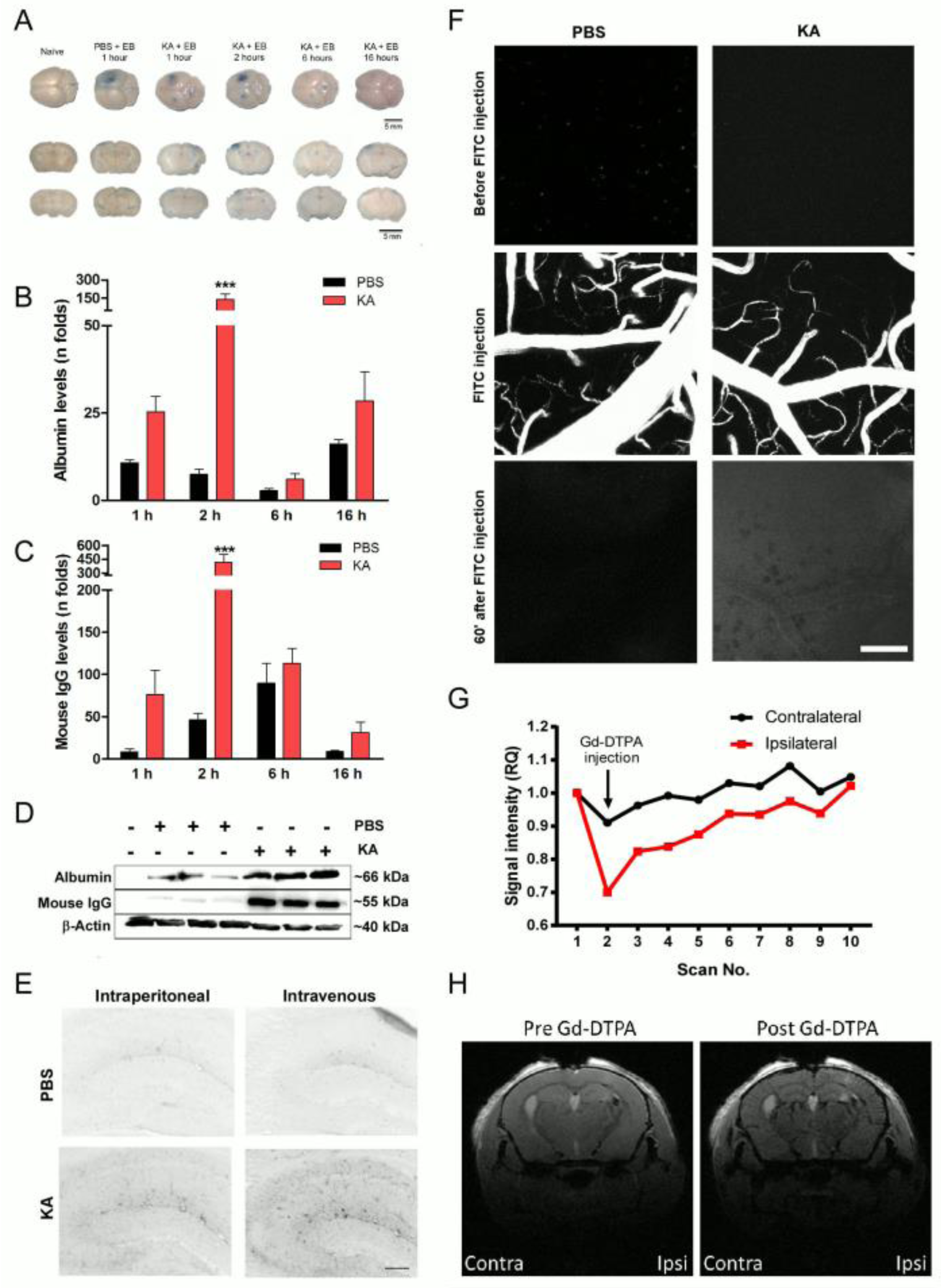
Blood-brain barrier (BBB) disruption after KA-induced status epilepticus in mice. (A) Extravasation of Evans Blue (EB) after IV injection after KA status epilepticus in mice. Scale bar, 5 mm. (B-D) Graphs and representative immunoblots (D) show protein levels (increase relative to naïve mice) of albumin (B) or IgG (C) after KA or PBS injection. Two-way ANOVA ***P<0.001. (E) Qualitative microscopic analysis showing extravasation of FITC-dextran at 2 hours after status epilepticus (from *n* = 3/group). Scale bar, 100 µm. (F) BBB disruption after status epilepticus. Two-photon images at 5 min before and 60 min after systemic injection of a 10 kDa FITC-dextran. Note increased intensity 1 hour after FITC injection in status epilepticus mice. Scale bar, 100 µm. (G-H) Contrast-enhanced MRI analysis of clearance rates of Gd-DTPA (742 daltons) in hippocampal regions after tail-vein injection of the contrast agent, 2 hours after status epilepticus. (G) Graph showing changes in Gd-DTPA clearance between contralateral (contra) and ipsilateral (ipsi) hemispheres (*n* = 10 for scan replications pre- and post-injection in the same mouse). (H) Representative MRI images depicting ipsi and contra brain hemispheres pre- and post-Gd-DTPA injection.

To determine whether BBB disruption at two hours was sufficient to allow passage of a macromolecule the size of Ant-134 (∼6 kDa), we performed IV or intraperitoneal (IP) injections of a fluorescein isothiocyanate (FITC)-conjugated 10 kDa dextran tracer 2 hours after status epilepticus. Brain slices from mice were then analysed by fluorescence microscopy for extravasation of FITC around cerebral microvessels. Both IV and IP injected FITC-dextran was observed outside microvessels and within brain parenchyma in mice subject to status epilepticus but not in vehicle-injected controls (Fig. 1E). This included the ipsilateral hippocampus, KA injection site and portions of the neocortex (Fig. 1E and data not shown). To extend these findings, we used live two-photon imaging through a cranial window to track FITC-dextran (10 kDa) extravasation after IV injection around cerebral microvessels after status epilepticus in living mice (Fig. 1F; fig. S1C). Beginning 1 hour after status epilepticus, imaging confirmed FITC signal outside small vessels within the neocortex ipsilateral to the side of seizure induction, which was not observed in non-seizure, vehicle-injected mice (Fig. 1F). Limitations on imaging depth prevented analysis of the extravasation within the hippocampus.

Finally, we imaged extravasation of magnetic resonance imaging (MRI) contrast agent (*25*) in mice 2 hours after status epilepticus. Using 7-T scanning, we observed enhanced contrast (manifested by an apparent signal disruption based on a T1 Flash MRI sequence) within the seizure-damaged hippocampus after systemic gadolinium (Dg-DTPA-800 Da) injection (Fig. 1G-H). While signal intensity returned to pre-gadolinium levels thereafter, this was delayed on the ipsilateral relative to the contralateral side, reflecting leakage of the gadolinium into surrounding brain tissue (Fig. 1H). Therefore, intraamygdala KA-induced status epilepticus in mice leads to focal disruption of the BBB sufficient to allow passage of a systemically-injected macromolecule such as an antagomir.

### Systemic delivery of Ant-134 timed with BBB disruption blocks miR-134 upregulation

We used locked nucleic acid (LNA)-modified cholesterol-tagged antagomirs to achieve potent and specific silencing of miR-134 (*11*). The targeting sequence and a 3D rendering of the predicted structure of Ant-134 and Ant-134 bound to miR-134 are shown in Figs 2A-C. Two hours after status epilepticus, mice were injected either IV or IP with 10 or 30 mg.kg^-1^ Ant-134, with dosing guided by previous studies (*17*). Control mice were systemically injected with a scrambled LNA-modified OGN (Scr). Brains were removed 24 hours later for analysis of miR-134 levels.

**Figure 2.**
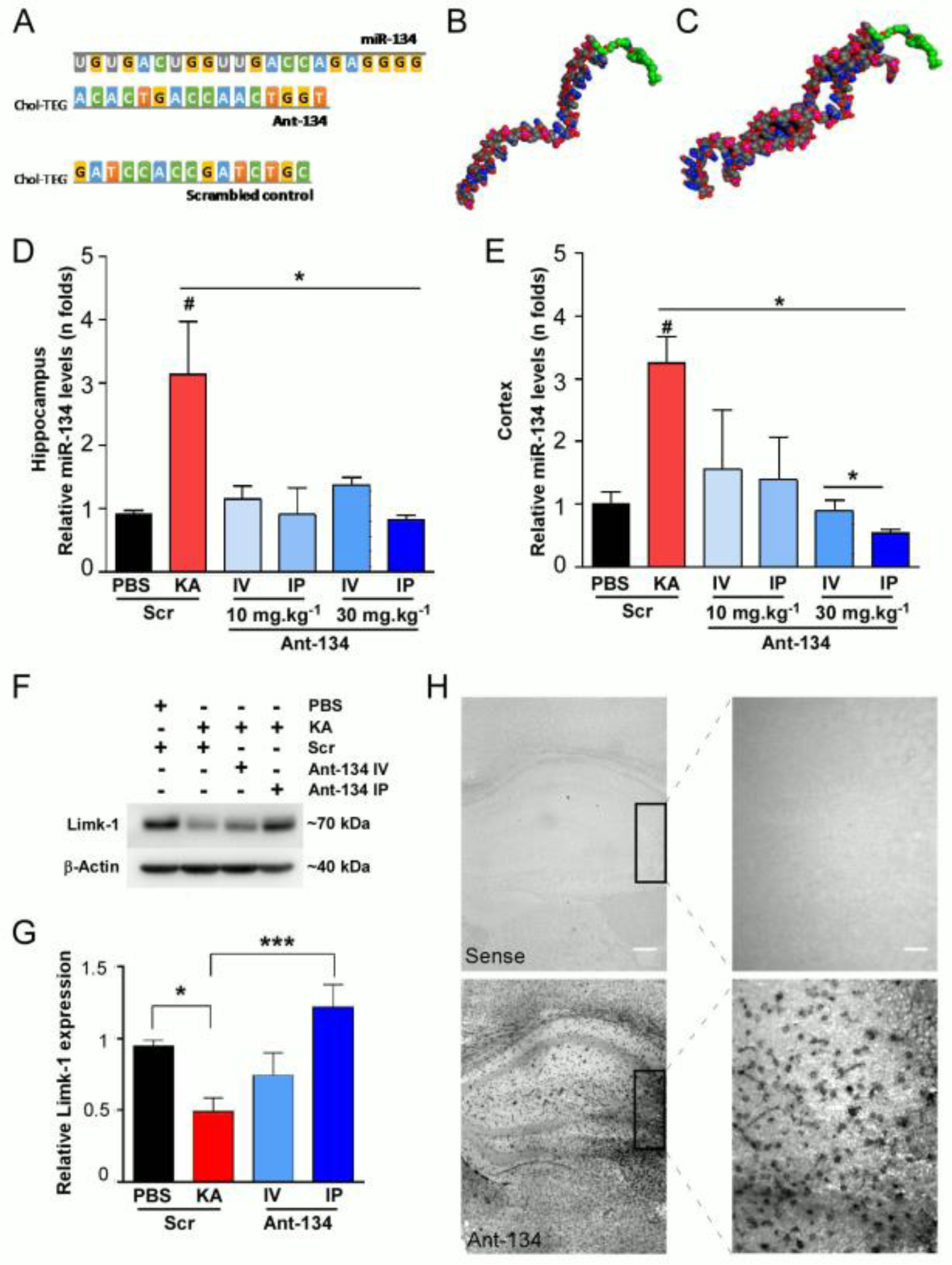
Seizure-facilitated entry of Ant-134 into the mouse brain after systemic injection. (A) OGN sequences of miR-134, Ant-134 and Scr. (B) Predicted model of Ant-134. Carbon atoms are depicted in grey, oxygen in red, nitrogen atoms blue, phosphorous atoms in pink. The carbons in the Col-TEG tag are coloured green. (C) Model of the predicted dimer of Ant-134 with miR-134. (D-E) Levels of miR-134 in the ipsilateral hippocampus (D) and cortex (E) of mice that were systemically injected with Ant-134 or Scr (10 or 30 mg.kg^-1^; IP or IV) at 2 hours after status epilepticus compared to PBS-Scr control mice. ANOVA *P<0.05 (*n* = 3-6/group). (F-G) Ant-134 (30 mg.kg^-1^) systemically administered by IP injection prevented the expected reduction of Limk1 protein levels induced by status epilepticus. Data were normalized to β-actin. ANOVA *P<0.05, ***P<0.001. (H) Representative images depict the presence of Ant-134 in the mouse brain 24 hours after IP injection (30 mg.kg^-1^). Sense (Scr inhibitor probe) was used as negative control. Scale bars represent 200 and 25 µm for magnifications 5x (left panel) and 40× (right panel), respectively.

Hippocampal (Fig. 2D) and neocortical (Fig. 2E) levels of miR-134 were ∼3 fold higher in mice subject to status epilepticus compared to vehicle controls, consistent with previous reports (*11*). IV and IP injection of either 10 or 30 mg.kg^-1^ Ant-134 blocked the upregulation of miR-134 after status epilepticus (Fig. 2D-E). We next measured levels of the miR-134 target Limk1 in brain samples from the mice. Consistent with previous reports (*11*), protein levels of Limk1 were lower in hippocampal samples from mice subject to status epilepticus, and this was partly reversed in Ant-134-injected seizure mice (Fig. 2F, G). Notably, target de-repression was greatest in mice given 30 mg.kg^-1^ IP injection of Ant-134 (Fig. 2F, G). Based on these findings and the less invasive delivery, an IP route was selected for studies thereafter. Finally, to confirm that systemically delivered Ant-134 after status epilpeticus reached the target tissue, we performed *in situ* hybridization (ISH) using specific probes for Ant-134 on mouse brain tissue sections collected 24 hours after injection. ISH confirmed strong Ant-134 staining within the ipsilateral hippocampus of mice that received an IP injection of Ant-134 (30 mg.kg^-1^) 2 hours after status epilepticus (Fig. 2H). The intensity of staining was particularly notable around the CA3 subfield, which is the main lesion site and source of SRS in the model (*26*).

### Systemic Ant-134 produces potent and lasting suppression of epilepsy in mice

Next, we equipped mice with implantable EEG telemetry units to assess whether systemically-delivered Ant-134 affected epilepsy development. Animals were then subjected to status epilepticus and, 2 hours later, given an IP injection of either 30 mg.kg^-1^ Ant-134 or Scr. Because clinical trials do not use non-targeting OGNs as the placebo arm, we verified that the basic epilepsy phenotype in Scr mice was similar to vehicle (saline) (fig. S2A-B). EEG recording units were activated during status epilepticus and for the next 2 weeks. They were then turned off and re-activated for one week of recording at 1, 2 and 3 months after status epilepticus to assess the permanency of any effects, after which mice were killed and brains processed for histology (see schematic in Fig. 3A).

**Figure 3.**
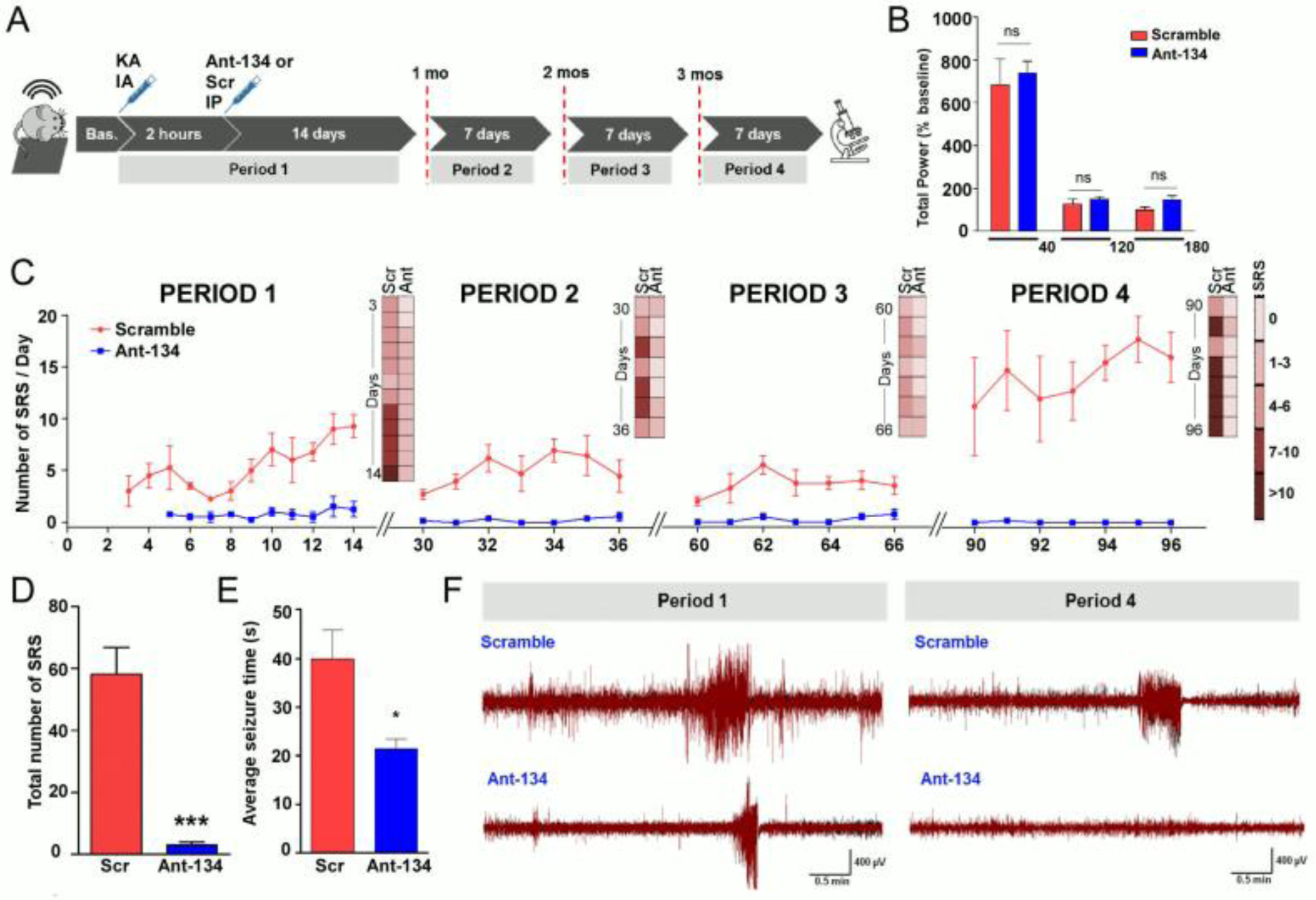
Potent and lasting anti-seizure effects of systemic Ant-134 in mice. (A) Experimental design for long-term epilepsy monitoring studies. After baseline (Bas.) EEG recording, status epilepticus was induced then 2 hours later mice were injected with Ant-134 or Scr control (IP, 30 mg.kg^-1^) and followed for 14 days (“Period 1”) and again for 7 days at 1 month (“Period 2”), 2 months (“Period 3”) and 3 months (“Period 4”). (B) No differences in the severity of the status epilepticus prior and soon after Ant-134 or Scr injections. X-axis plots time (in min) after status epilepticus between KA and lorazepam injections; lorazepam and Ant-134/Scr injection; and up to 1 hour after Ant-134/Scr injection. (C) Spontaneous recurrent seizures (SRS) per day per treatment group over time depicting the total number of seizures per group per EEG activation period. (D) Graph showing overall reduction of SRS (*t*-test; P< 0.001). (E) Duration of SRS in Ant-134 compared to Scr mice (*t*-test; P = 0.042). (F) Representative EEG traces depict SRS at the last day of recording for “Period 1” (left) and “Period 4” (right) for Scramble-(top) and Ant-134-treated mice (bottom). Note, Ant-134 (“Period 4”) EEG trace is similar to baseline recordings. <u>Data was from n = 5 (Scr) and 6 (Ant-134) (Period 1), with final Period data from n = 4 (Scr) and 5 (Ant-134).</u>

Systemic injection of Ant-134 did not alter epileptiform EEG parameters during the first 3 hours after status epilepticus, confirming the initial precipitating injury was equivalent between groups that would be subsequently monitored (Fig. 3B). Seizure monitoring in freely-moving Scr-treated mice during the first recording period determined that SRS emerged within 3 - 4 days of status epilepticus, consistent with the known course of epilepsy in the model (*11*). SRS occurred daily in most Scr mice, ranging from 1 - 20 per day. In contrast, mice given a systemic injection of Ant-134 after status epilepticus displayed only rare seizures during the first two-week recording period and these emerged slightly later (at 5 - 7 days) (see “Period 1” from Fig. 3C and 3F; fig. S2C). In the first 2 weeks of monitoring, we observed ∼85% fewer SRS in Ant-134-injected mice compared to controls (“Period 1”, Table S1; fig. S2A). A secondary measure of seizure burden, based on the duration of seizures, revealed ∼92% reduction in Ant-134 injected mice when compared with Scr-injected mice (fig. S2B).

Ant-134-injected mice continued to display dramatically lower rates of SRS compared to Scr mice during monitoring at 1, 2 and 3 months after status epilepticus (see “Periods 2, 3 and 4”, respectively, Fig. 3C and 3F; fig. S2C) and presented an overall average of 6.14 (out of 7) days seizure-free (see “Periods 2 to 4”, Table S1). Most notably, during the final week of seizure monitoring, only a single SRS was detected in one of the Ant-134 mice compared to an average of ∼100 epileptic seizures in the Scr group (“Period 4”, Table S1). This was equivalent to a 99.5% lower seizure rate in Ant-134 mice in the third month, and around 93.4% over all periods (Fig. 3D). On average, the duration of SRS was ∼45% shorter in Ant-134-than in Scr-injected mice over all monitoring periods (Fig. 3E).

### Histological findings in Ant-134-treated mice

Brains from mice obtained at the end of EEG monitoring (*i*.*e*. three months after status epilepticus) displayed modest hippocampal atrophy and were macroscopically similar between Ant-134 and Scr-injected mice (Figure 4). However, tissue sections from Ant-134-treated mice displayed lower glial fibrillary acid protein (GFAP) staining when compared to Scr, suggestive of reduced astrogliosis (Fig. 4A; fig. S3A). Counts of Iba1-positive cells, likely microglia, were not different between the groups (Fig. 4B; fig. S3B). Gross neuron loss was similar between groups, as assessed using a semi-quantitative scoring system, but Ant-134-treated mice had higher numbers of inhibitory parvalbumin interneurons (*27*) (Fig. 4C; fig. S3C).

**Figure 4.**
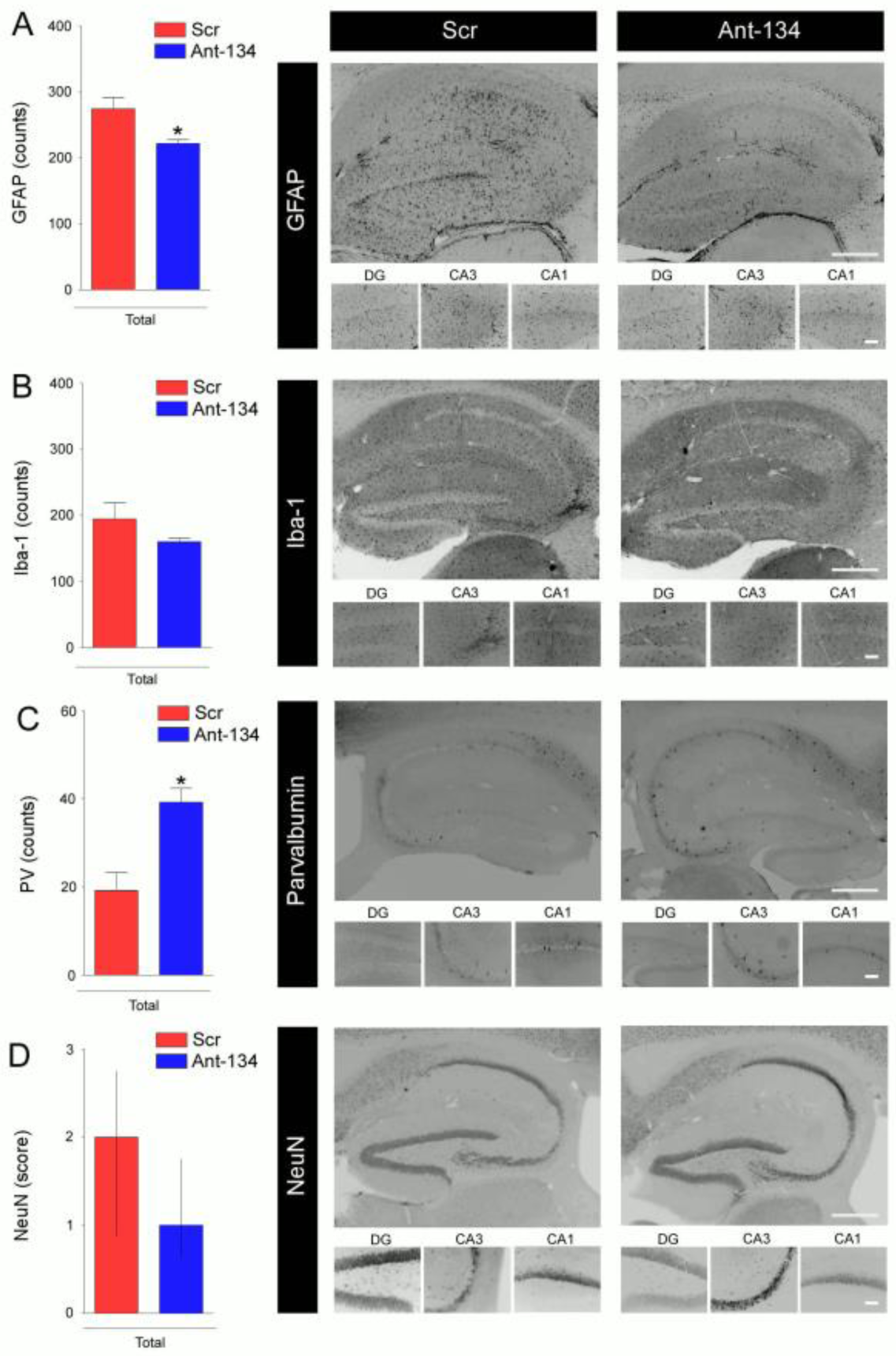
Histological findings in Ant-134-treated mice. Graphs and representative images of brains obtained at the end of long-term epilepsy monitoring stained for (A) GFAP (astrocytic marker), (B) Iba1 (microglia), (C) Parvalbumin (PV; a marker for GABAergic interneurons) and (D) NeuN (broad neuronal marker). Representative photomicrographs of each cellular marker show the whole ipsilateral hippocampus (5X; top) and hippocampal sub-regions (DG, CA3 and CA1; 20×; bottom) for Scr- and Ant-134 treated mice (P<0.05 compared to Scr). <u>n = 4 (Scr), n = 5 (Ant-134).</u> *P < 0.05 compared to Scr.

We used additional brain tissue sections from the post-monitoring mice to stain for Ant-134. The absence of the antagomir at the end of epilepsy monitoring would support a disease-modifying effect. Ant-134 staining was readily observed in positive control samples obtained 24 hours after status epilepticus from mice injected with Ant-134 whereas Ant-134 staining was not detectable in the brain tissue sections from mice at the end of epilepsy monitoring (fig. S3D). Therefore, the lack of SRS in Ant-134 mice was unlikely to be due to the continuing presence of Ant-134 within the brain.

### Anti-seizure mechanism of Ant-134

We next sought to explore the mechanism(s) by which Ant-134 produces anti-seizure effects. Limk1 has a direct role in the control of dendritic structure and function function (*14*) and Ant-134 restores Limk1 levels after status epilepticus (*11*) (and see Fig. 2F-G). We therefore hypothesised that the seizure-suppressive effects of Ant-134 may be due in part to de-repression of Limk1. Accordingly, lowering Limk1 levels should revert the anti-seizure phenotype. To target Limk1, we used LNA-GapmeR OGNs (Gap-LIMK) which deplete target mRNAs via an RNase H-dependent mechanism. We first performed *in vitro* screening to identify an optimal sequence that reduced Limk1 levels by an amount equivalent to that found to be recovered by Ant-134 treatment (fig. S4A-C). The ability of an ICV injection of Gap-LIMK to knockdown Limk1 was then confirmed *in vivo* (Fig. 5B). Next, we equipped mice for EEG telemetry recordings. One group of mice then received ICV injections of Gap-LIMK while the other group received a scrambled (Scr) version and all mice were injected with Ant-134 (30 mg.kg^-1^ IP) 2 hours after status epilepticus. Gapmer injections were repeated every second day. SRS frequency was quantified as before for a period of 2 weeks (see schematic in Fig. 5A). Note, injection of GAP-LIMK alone did not produce epilepsy in mice (data not shown).

**Figure 5.**
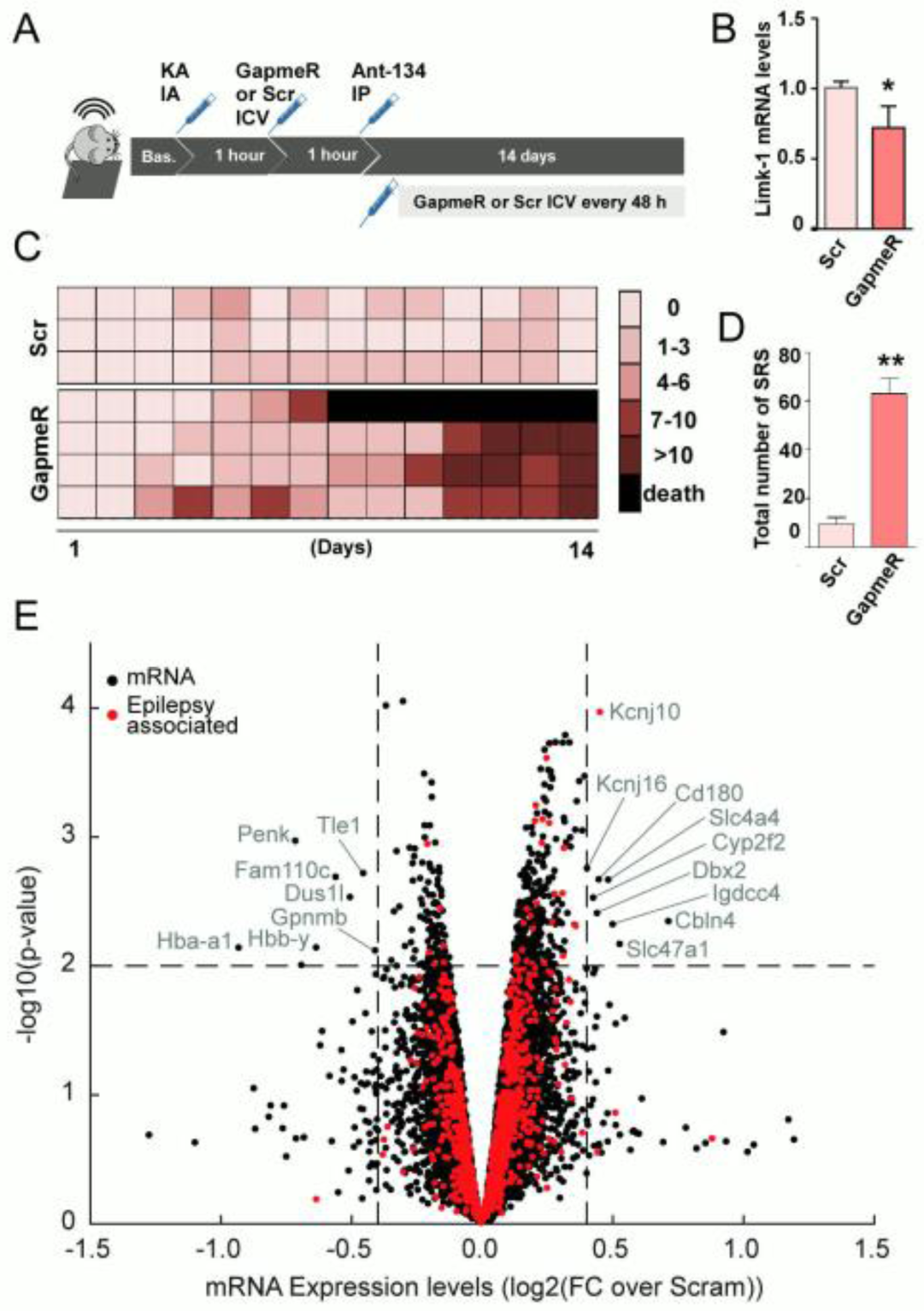
Molecular effects and mechanism of Ant-134. (A) Mice received ICV injection of Gapmer to reduce Limk1 (or Scr) 1 hour after status epilepticus and then injection of Ant-134 (IP, 30 mg.kg^-1^; at 2 hours) followed by 2 weeks of EEG recording. (B) Reduction of *Limk1* mRNA in the hippocampus 24 hours after a single Gapmer injection in comparison with Scr control (*n* = 3/group; t-test; **P = 0.017). (C) Individual seizure counts per day for Ant-134 mice that received either Gapmer-Limk or Scr. Each line represents data from the same mouse over time, and each row represents the seizure counts for the respective day. (D) Total number of SRS over 2 weeks of EEG recordings comparing [Ant-134+(Scr)] mice to [Ant-134+(GapmeR)] mice (*n* = 3/group; mouse that died was excluded for statistical purposes; t-test; **P=0.013). (E) Microarray analysis showing gene expression changes 24 hours after status epilepticus in Ant-134-treated mice compared to Scr (*n* = 3/group). Note, select differential expression induced by Ant-134. Dashed lines are drawn at log2FC = 0.4 and −0.4 (equivalent to +/-1.3 fold change) and –log10(p-value) = 2.0 (equivalent to FDR p-value = 0.01). mRNA with log10(p-value) > 2.0 and log2FC >0.4 or < −0.4 are labelled, and those identified as associated with epilepsy are marked in red.

As expected, the frequency of SRS was very low in mice given systemic Ant-134 after status epilepticus and ICV injections of the non-targeting Gapmer (Fig. 5C-D). The frequency of SRS in Ant-134-treated status epilepticus mice given Gap-LIMK was initially similar but within a few days increased, reaching a frequency similar to the basic model (i.e. average of 4 - 6 SRS per day) (Fig. 5C-D). Thus, reducing Limk1 levels *in vivo* partly obviates the anti-seizure phenotype of Ant-134, supporting de-repression of Limk1 in the anti-seizure mechanism of Ant-134.

Based on these findings and the known multi-targeting effects of miRNAs, we sought to identify other potential targets altered by Ant-134, performing transcriptional profiling of the hippocampus of mice 24 hours after status epilepticus. This identified 16 differentially expressed mRNAs between Scr and Ant-134-treated mice (Figure 5E). This suggests that the downstream effects of Ant-134 at the transcriptome-level during epileptogenesis may be subtle, complex, and/or time-sensitive. Nevertheless, significantly dysregulated mRNAs included plausible candidate genes for an anti-seizure mechanism, including increased levels of *Kcnj10*, an inward-rectifying potassium channel subunits, the loss of which may cause epilepsy in humans (*28*).

### Neuronal localisation of miR-134 in human pharmacoresistant epilepsy

Although previous studies reported that the level of miR-134 was increased in brain samples from patients with pharmacoresistant temporal lobe epilepsy (*11, 12*), its cellular expression in human brain is poorly understood. We therefore used ISH to identify the cell types expressing miR-134 in human brain. Autopsy control and hippocampal tissue sections from patients with pharmacoresistant temporal lobe epilepsy both displayed miR-134 staining and this appeared exclusively neuronal (Figure 6A and fig. S5A). To confirm the cell phenotype expressing miR-134 in human brain we performed fluorescence ISH using markers for neurons (NeuN; Figure 6B) and astrocytes (GFAP; fig. S5B). This confirmed miR-134 is restricted to neurons in the human hippocampus, suggesting neurons are a key target of Ant-134.

**Figure 6.**
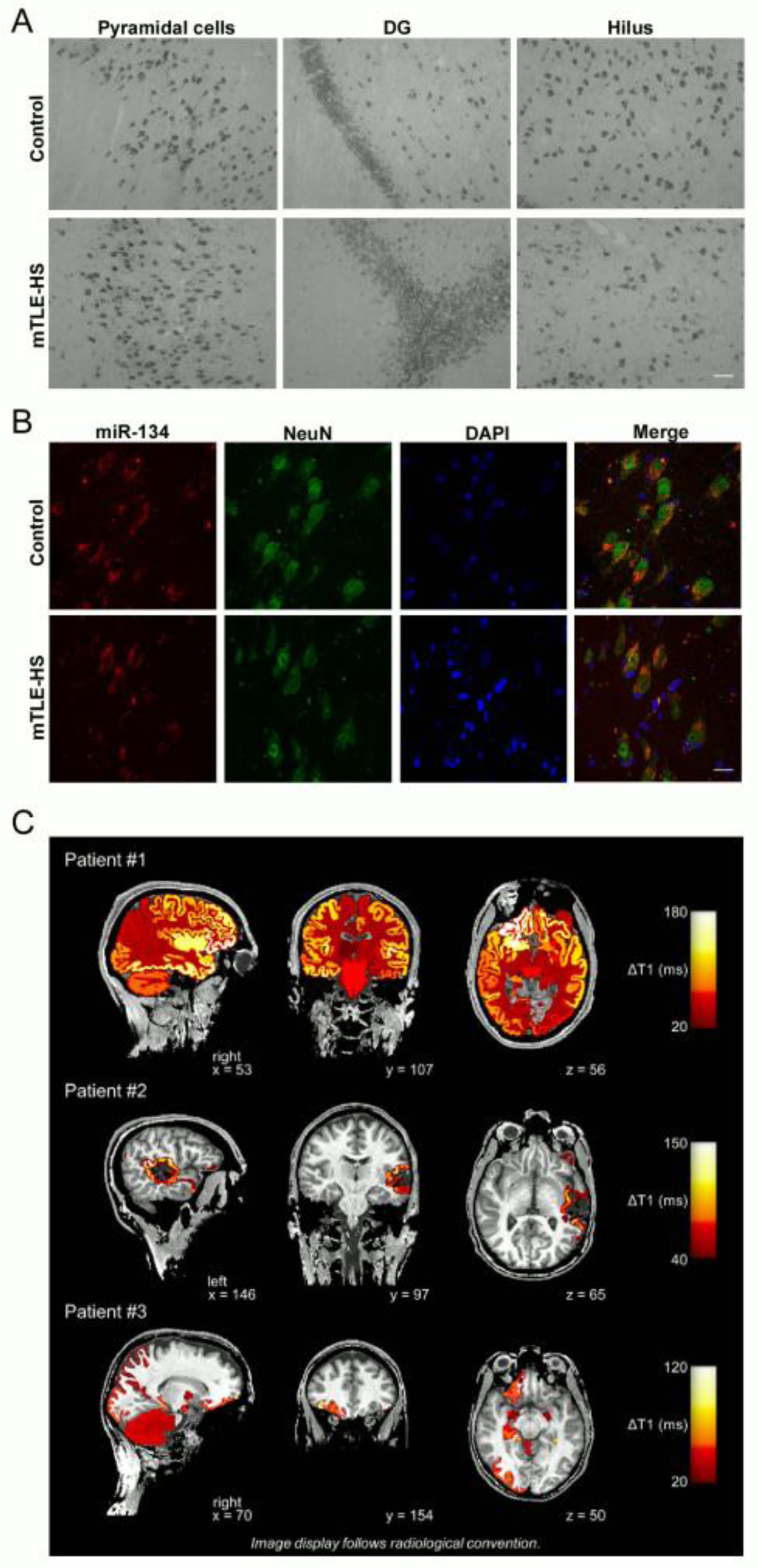
Neuronal expression of miR-134 in human brain and BBB disruption after seizures in human epilepsy. (A) Representative ISH images showing miR-134 expression in the hippocampus from temporal lobe epilepsy patients and autopsy controls. Scale bar, 100 µm. (B) FISH showing miR-134 co-localizes with the neuronal marker NeuN. Scale bar, 50 µm. (C) BBB dysfunction after seizures in epilepsy patients. ΔT1 indicating postictal enhancement of gadolinium-based contrast agent in three patients. ΔT1 computed by subtracting postictal qT1 from interictal qT1. Higher ΔT1 values, thus, reflect a greater postictal enhancement of contrast agent. Patient #1: male, 24 years old, seizure duration: 1.14 min, seizure-onset-to-injection time interval: 17 sec, injection-to-MRI time interval: 25 min, patient showed secondarily generalized tonic-clonic seizure, seizure semiology: sleep -> arousal -> sign of 4 with left arm extension and right arm flexion -> bilateral tonic-clonic phase, non-lesional MRI, ictal EEG onset: right frontal, presumed seizure onset zone: right frontal. Patient #2: male, 34 years old, seizure duration: 57 sec, seizure-onset-to-injection time interval: 12 sec, injection-to-MRI time interval: 26 min, patient showed a focal seizure, seizure semiology: HV -> oral automatisms, behavioral arrest -> manual automatisms, MRI: resected cavernoma in left temporal lobe, ictal EEG onset: left temporal, presumed seizure onset zone: left temporal. Patient #3: female, 34 years old, seizure duration: 26 sec, seizure-onset-to-injection time interval: 9 sec, injection-to-MRI time interval: 32 min, patient showed a focal seizure, seizure semiology: sleep -> arousal -> opens eyes -> non-forced head devotion to the right side -> oral automatisms -> drowsy state, MRI showed ictal EEG onset: right frontal, presumed SOZ: right frontal implantation of electrodes in right olfactory sulcus, resection of lesion (histology: low-grad, hamartoma-like lesion), patient is postoperatively seizure-free (Engel 1a according to (*40*)).

### BBB impairment following seizures in humans

Human MRI studies using contrast agents have shown that epilepsy-causing brain injuries are associated with temporal and anatomical BBB dysfunction (*21*), and BBB disruption can occur after seizures (*29*). To provide a clinical context to our animal model findings, we analysed contrast-enhanced MRI brain imaging studies of a set of three patients who recently experienced seizures (Fig. 6C). Two patients experienced a focal seizure while one patient had a secondary generalized tonic-clonic seizure. ΔT1 indicated postictal enhancement of gadolinium-based contrast agent 25 to 32 min after seizure onset and injection of contrast agent. The postictal enhancement was anatomically associated with the spread of ictal activity as determined by clinical observation and results of other diagnostic modalities (Fig. 6C). These findings provide evidence for peri-ictal disruption of the BBB occurring after seizures in humans.

## Discussion

Anti-epileptic drugs fail to control seizures in one third of patients and do not have disease-modifying effects. The targeting of miRNAs has emerged as a promising therapeutic approach but the clinical deployment of miRNA-blocking OGNs is limited by the need to circumvent the BBB. The present study demonstrates that, when timed with BBB disruption after status epilepticus, a single systemic injection of an antisense miRNA inhibitor produces potent and lasting suppression of epilepsy in mice. These findings expand the evidence for the disease-modifying potential of antagomirs targeting miR-134 and offer a clinically-viable route for delivery of this promising experimental therapy for epilepsy.

Targeting miR-134 using OGNs has been particularly notable for the potency and disease-modifying actions demonstrated across different *in vivo* animal models (*11-13, 18*). Delivering OGNs to the brain is a key challenge however, as is knowledge of the mechanism of action because the multi-targeting actions of miRNAs raise unique safety concerns (*6*). Currently, OGN delivery for brain diseases is expected to require an intrathecal or ICV route, or some form of conjugation or encapsulation technique. However, BBB disruption accompanies many epilepsy-triggering brain injuries (*21*), potentially facilitating non-invasive (i.e. systemic) delivery. The present study addressed both delivery and mechanism of action, revealing that systemic injection of Ant-134 can prevent SRS in experimental epilepsy, principally through a single target.

While prolonged or repeated seizures are known to cause disruption of the BBB, the focus until now has been on understanding how this contributes to epilepsy (*20, 29*). Here we asked whether BBB disruption after status epilepticus could be appropriated to deliver a systemically-administered antagomir to treat epilepsy. Using multiple assessments of BBB integrity, including extravasation of plasma markers and live brain imaging, we identified a disruption 2 hours after status epilepticus sufficient to allow passage of systemically administered Ant-134 into the brain parenchyma. While the upper size limit of this disruption was not pursued, the selectivity size and timing align with earlier histological and biochemical findings in the same model (*24*) and results in other models of status epilepticus (*20*). We focused on IV and IP routes for delivering Ant-134 to the brain although other minimally invasive routes (e.g. subcutaneous) and either earlier or later time-points for injection may also be effective. Target engagement after systemic injection was confirmed by measurement of miR-134 knockdown and increased Limk1 and demonstrated by staining for the antagomir in the damaged CA3 subfield, a key site of ictogenesis during SRS in the model (*26*). However, delivery to other sites may also be important (*4*). Indeed, BBB disruption occurred elsewhere in the model and we observed knockdown of miR-134 by Ant-134 in the neocortex after systemic delivery.

Our data from long-term epilepsy monitoring revealed remarkable disease-modifying effects of silencing miR-134 using systemically-delivered Ant-134 after status epilepticus. Seizures were reduced by 99.5% during the final week of monitoring at three months. This exceeds the performance of other anti-epileptogenic approaches currently under investigation (*5*). The anti-seizure effect was also somewhat superior to the outcome when Ant-134 was delivered by ICV route in the same model, perhaps due to differences in the amount of Ant-134 that reached brain sites (*11, 13*), including the contralateral hippocampus (*23*), or the timing of Ant-134 delivery, which was slightly later in the present study (*11, 13*). Appraisal of how these individual variables affect performance may yield further improvements in effectiveness. Further studies will be needed to determine if Ant-134 can prevent SRS following epileptogenic insults besides status epilepticus (*21, 22*).

The multi-targeting nature of miRNAs makes understanding the mechanism(s) by which their inhibition produces anti-seizure effects uniquely challenging (*30*). Past studies showed that miR-134 targets Limk1, a protein that promotes dendritic spine expansion by regulating actin dynamics through phosphorylation of cofilin (*14, 18*). Here we used Gapmer OGNs to produce a partial reduction in Limk1 levels, an approach that is physiologically relevant to the scale of protein suppression by a miRNA on a target (*7, 14*). This was sufficient to increase SRS in Ant-134-treated epileptic mice, providing *in vivo* support for the disease-modifying effects being a result of de-repression of Limk1. This implies the anti-seizure effects of Ant-134 result from adjustment to synaptic microstructure, perhaps curtailing the capacity of neuronal networks to propagate hypersynchronous discharges (*31*). The finding fits with an emerging view that some miRNAs produce strong effects on single targets, perhaps because of spatial co-localisation (*32*). Notably, other anti-seizure antagomirs have been reported to derive effects from de-repression of single ion channels (*30*). Nevertheless, Limk1 inhibition did not fully obviate the anti-seizure effects of Ant-134 suggesting other targets of miR-134 may contribute to the observed phenotype or pathophysiology. On this note, we observed reduced astrogliosis and sparing of interneurons in brain sections from Ant-134 mice, both of which could account for reduced seizures (*27, 33*), but could also be a consequence of fewer seizures. Our profiling of the hippocampal transcriptional landscape in Ant-134-treated mice identified further potential targets that might be explored including potassium channels. Studies in human cells or tissues will also be needed to assure clinical relevance of the Ant-134 mechanism and target pool.

The present findings improve the prospects for the development of an antagomir therapy for epilepsy. An antagomir therapy has already completed clinical trials for hepatitis C (*34*), various others are in development, and OGN-based treatments are in clinical use for neuromuscular diseases (*1*). Although pK and toxicity studies have not yet been performed, broad safety might be assumed on the basis that mice in our study were given a systemic injection of Ant-134 and lived for three months. This may be helped by the largely CNS expression of miR-134 (*14*), and we show here it is specifically expressed by neurons in the human brain. While manipulation of a miRNA that regulates dendritic spines carries risks, Ant-134 injections in mice (*11, 18*) and rats (*31*) were not reported to impact hippocampal function or animal behaviour.

How might an Ant-134 therapy be tested in patients? The closest clinical scenario to the studies here would be deployment of the antagomir to prevent the development of epilepsy in at-risk individuals. Such a trial would require, however, the identification of biomarkers of human epileptogenesis (*5*). Although validated biomarkers do not yet exist, increased miR-134 levels in biofluids including plasma have been reported in some epilepsy patients (*35*) and the co-development of devices for point-of-care miRNA testing could further enable translation (*36*). However, preclinical studies show that Ant-134 can prevent SRS in models in which miR-134 is not dysregulated (*12*). A first clinical trial could instead focus on testing in pharmacoresistant focal epilepsy. An injection might be offered to patients awaiting neurosurgery such that the target tissue would be excised later and studied for antagomir effects, as proposed for gene therapy trials for this patient group (*37*). Last, although we took advantage of the BBB disruption that accompanies prolonged seizures, other methods for artificially opening the BBB may also facilitate delivery of Ant-134 in patients such as hyper-osmotic treatments or ultrasound.

In summary, the present study shows that timed systemic delivery of Ant-134 produces potent and long-lasting disease-modifying effects which are partly mediated by Limk1. These findings suggest that this miRNA-based therapy can be delivered using a clinically-relevant route, and provides the rationale for pursuing further pre-clinical and clinical studies, which are needed to evaluate the safety and efficacy of this potential candidate for the treatment of epilepsy.

## Material and Methods

### Study design

Treatment (Ant-134) and control (Saline or Scramble) groups were randomly assigned before the induction of *status epilepticus*, and the researcher was blind to the experimental groups during injections. EEGs were randomly assigned for analysis, which was blindly performed by two individuals. The sample size was defined based on previous work from our group(*11*), revised and approved by biostatistician at Royal College of Surgeons in Ireland (RCSI), and not altered during the course of the study. For the EEG long-term recordings, the selected endpoint was the evaluation of epilepsy development in mice treated with Ant-134 in comparison with controls. The EEG samples were collected for 2 weeks after *status epilepticus* and for one week at later time points (exactly at 1, 2 and 3 months after *status epilepticus*). This data collection schedule was defined considering experimental limitations (battery life of telemetric EEG devices). No outliers or data were excluded in this study, and statistical correction was applied for animals that died during the EEG long-term monitoring or did not present seizures at certain days (see Statistical Analysis).

### Animals

All animal experiments were performed in accordance with the European Communities Council Directive (2010/63/EU), the NIH Guide for the Care and Use of Laboratory Animals, and followed ARRIVE guidelines. Procedures in mice were approved by the Research Ethics Committee of the Royal College of Surgeons in Ireland (REC-842), under license from the Health Products Regulatory Authority (AE19127/001), Dublin, Ireland. Adult male C57BL/6 mice (∼25 g; 6-7 weeks old) obtained from RCSI’ s biomedical research facility (original stock from Harlan, Oxon, Bicester, U.K.), were used in all studies. Animals were housed (up to 5 mice per cage) in on-site barrier-controlled facilities having a 12 h light-dark cycle with *ad libitum* access to food and water. Single housing was used for long-term EEG experiments.

### Seizure model and acute EEG assessments

Mice were anesthetized with isoflurane (isoflurane; 5% induction, 1–2% maintenance) and placed in a mouse-adapted stereotaxic frame. Body temperature was maintained within the normal physiological range with a feedback-controlled heat pad (Harvard Apparatus, Kent, UK; Holliston, MA). For acute assessments (e.g. BBB study and molecular assessments after Ant-134 injections), after making a midline scalp incision, the Bregma was located, and three partial craniectomies were performed for the placement of skull-mounted recording screws (Bilaney Consultants). A fourth craniectomy was drilled for the placement of a guide cannula (coordinates from the Bregma: anterior-posterior (AP) = – 0.94 mm, lateral (L) = – 2.85 mm). The cannula and electrode assembly was fixed in place with dental cement, and the mouse was placed in a heated chamber for recovery prior to further studies. Mice were then placed in an open Perspex box and connected to a Xltek® EEG brain monitor amplifier (Natus). After baseline EEG was established, status epilepticus was triggered in freely-moving awake mice by intraamygdala microinjection of KA (0.3 µg in 0.2 µl of PBS) (Sigma-Aldrich, Ireland), as previously described (*11*). Control mice were injected with the same volume of PBS (pH adjusted to 7.4). After 40 min, all mice, including controls, received lorazepam (8 mg/kg; 10 ml/kg; IP) to curtail seizures and reduce morbidity and mortality. Mice were recorded for 1-3 h thereafter, according to experimental design, before being disconnected and placed in a warmed recovery chamber.

Mouse EEG data were analyzed and quantified using LabChart 8 software (ADInstruments, Oxford, U.K.). Seizures were defined as high-amplitude (> 2 x baseline) high-frequency (> 5 Hz) polyspike discharges lasting > 5 sec. Status epilepticus was defined as at least 5 min of continuous seizure activity. From the EEG recordings we calculated the total power and % of total power before lorazepam injection, as previously described (*11*). EEG total power was plotted as percentage of baseline recording (each animal’s EEG power post seizure compared to its own baseline EEG).

#### Ant-134 and administration protocol

Ant-134 (anti-mmu-miR-134; MW: 6127.3 Da) or scramble (Scr) control (both from Exiqon, LNA-modified and 3’-cholesterol–modified oligonucleotides) in PBS (pH adjusted to 7.4) was prepared in a sterile solution concentration of 7.5 mg.ml^-1^, and injected IV or IP at 10 or 30 mg.kg^-1^ in mice at 2 h after intraamygdala KA-induced status epilepticus.

#### Long-term EEG telemetry recordings

Under isoflurane anaesthesia and using aseptic technique, mice were placed in a mouse-adapted stereotaxic frame. A mid-line incision was performed in the skin overlying the skull and Bregma located. Four partial and one complete craniotomy was performed. The leads from a DSI telemetry unit (Model: F20-EET, Data Systems International - DSI) were affixed to the skull for bi-lateral EEG recordings (Ponemah v6.30 software, DSI) and the unit placed in a subcutaneous pocket created in the upper back. A guide cannula was affixed (resting on the dura mater) for the intraamygdala injection, as before. The electrode-cannula assembly was fixed in place by dental cement and the mouse was removed from the stereotaxic frame and allowed to recover. Mice were randomly and blindly assigned to experimental groups (KA+ Saline, KA+Scr or KA+Ant134), returned to home cages and provided with soft food and fluids to aid recovery. Forty-eight hours later, EEG units were activated and a baseline recording was obtained before triggering status epilepticus as above followed by lorazepam after 40 min. Two hours later, mice were injected with saline, Scr or Ant-134 (30 mg.kg^-1^; IP; 10ml.kg^-1^). Continuous EEG data were then collected for 14 consecutive days (sampling epoch called “Period 1”). Telemetry units were then turned off and only turned on exactly 1 month later for one week of continuous EEG recording (sampling epoch called “Period 2”; days 30^th^ to 36^th^ after status epilepticus). Devices were deactivated again and re-activated at 2 and 3 months after status epilepticus (sampling epochs, respectively, “Period 3” / days 60^th^ to 66^th^; and “Period 4” / days 90^th^ to 96^th^). At the end of the entire experiment, mice were deeply anaesthetised and perfused with ice-cold saline and brains removed for immunohistochemistry and *in situ* hybridization analysis.

Telemetry EEG data were reviewed and manually scored by an observer unaware of the experimental treatment. The overall presence/number of spontaneous recurrent seizures (SRS) per day of recording and the average seizure time were quantified. Seizure burden was assessed only during “Period 1” to ensure statistical power was kept as initially planned. As before, epileptic seizures were defined as high-frequency (>5 Hz), high-amplitude (>2× baseline) polyspike discharges of ≥5 s duration. Quantified data was analysed as described below (Statistical Analysis).

#### In vivo Limk1 silencing by GapmeRs

Mice were equipped with telemetry devices, as described before. Schematic for the experimental design can be seen in Fig. 5B. Briefly, after a baseline EEG recording, mice underwent intra-amygdala KA-induced SE, randomly/blindly assigned in groups, and 1h later were intracerebroventricularly (ICV) injected with the first dose of GapmeR to silence Limk1 mRNA (called only “GapmeR”; n=4; 0.01 nmol/2µl) or its scramble negative control (called “Scr”; n=3; 0.01 nmol/2µl). One more hour after the ICV injection (2h after SE), all mice were systemically injected with Ant-134 (30 mg.kg^-1^, IP). Long-term EEG was continued for 2 weeks, and mice received ICV doses of GapmeR or Scr every 48h for the overall recording time (total of seven administrations on experimental days 1, 3, 5, 7, 9, 11 and 13). EEG data were reviewed and manually scored by an observer unaware of the experimental treatment. The overall presence and number of spontaneous recurrent seizures (SRS) per day of recording were quantified, as before.

#### Qualitative BBB extravasation by Evans Blue

At 1 h, 2 h, 6 h and 16 h after status epilepticus, mice (*n* = 3 /group) were intravenously (IV) injected with the azo dye 3% Evans Blue (5 ml/kg) to macroscopically and qualitatively assess BBB disruption. Five minutes after dye injection, mice were deeply anesthetized (pentobarbital) and transcardially perfused with ice-cold saline followed by 4% PFA, to clean blood components and fix the tissue, respectively. Images were obtained using a stereomicroscope from the whole brain and 1 mm coronal slices (dorsal and ventral hippocampus).

#### FITC-dextran extravasation

After acute surgical/EEG procedures as before, mice were intravenously (IV; tail vein; 5 ml/kg) or intraperitoneally (IP; 10 ml/kg) injected with the fluorescent reporter FITC-Dextran (10 kDa; 40 mg.kg^-1^) at 2 h after KA or PBS intraamygdala injection (*n* = 3/group). Five or 15 min after dye injection (for IV and IP, respectively), mice were deeply anesthetized (pentobarbital) and transcardially perfused with ice-cold saline followed by 4% PFA and whole brains snap frozen and sectioned (40 µm; coronal; CM1900 cryostat, Leica). Slices were protected from light, placed on a microscope slide and covered with a coverslip and qualitative assessed by imaging using imaged using a Hamamatsu Orca 285 camera attached to a Nikon 2000s epifluorescence microscope (Micron Optical).

#### Two-photon live imaging

For *in vivo* brain imaging, mice underwent a surgical procedure to create a cranial window. Briefly, anaesthetised mice were placed in a stereotaxic frame with controlled body temperature. The surgical procedure comprised of a 5 mm inner diameter craniotomy on the skull surface (over the somatosensory area between the Bregma and lambda) of the mice without disrupting the dura mater, and the placement of a clear sterile glass plate (fixed with acrylic cement) to cover the cranial window. Mice were also implanted with a guide cannula for intra-amygdala injection of PBS or KA prior to two photon imaging. After the procedure, mice were allowed to recover in a heated chamber and returned to their cages for a week. On the day of imaging, KA or PBS injection was performed as before. For imaging, the mouse was placed on an imaging stage under 1-2% isoflurane anaesthesia and Immersol W was placed between the W Plan-Apochromat 20× (NA 1.0) lens of the microscope and the window. Imaging took place over 2 h. During this time body temperature was controlled and maintained at 36.5– 37.5 °C using a heating pad connected to a controller and eyes were covered with eye protecting gel. After baseline images were captured, the fluorescent reporter FITC-Dextran (10 kDa) was IV injected as above, 1 h after intraamygdala KA or PBS injection. The live extravasation was recorded over time until the fluorescent reporter signal faded from vessels.

At 2 h after KA or PBS, mice were injected a second time with FITC-Dextran, as before, and the cumulative extravasation observed. Z-stack images were acquired every 60s using an upright Carl Zeiss LSM 710 NLO and Zen 2008. A Coherent Chameleon Vision II laser operating at 780nm with 40.0% laser power (64 Mw at sample) and a 500-550nm emission band pass filter was used for excitation and detection. After the procedure mice were euthanized. Maximum projection images were prepared using FIJI (*38*).

#### MRI in mice

BBB integrity was assessed 2 h following induction of status epilepticus (or intraamygdala PBS injection, controls), using a dedicated small rodent 7-T MRI system (www.neuroscience.tcd.ie/technologies/mri.php). Mice were anaesthetised with 5% isoflurane and placed on an MRI-compatible support cradle with a built-in system for maintaining the animal’s body temperature at 37 °C and for physiological monitoring (ECG and respiration). A cannula was flushed with saline and inserted into the tail vein. Accurate positioning was ensured by acquiring an initial rapid pilot image, which was then used to ensure the correct geometry for all subsequent scans. Compromises of the BBB were then visualised in high resolution T1 weighted MR images (resolution, 0.156 × 0.156 × 5 mm^3^; field of view: 20 × 20 × 17.9 mm^3^; matrix; 128 × 128 × 30; TR/TE: 500/2.7 ms; flip angle: 30o; number of averages: 3; acquisition time: 2 min, 24 sec; Repetitions: 12) before and after tail-vein injection of 100 µl of a 1 in 3 dilution of Gd-DTPA (0.5 M stock solution, Bayer). MRI analysis was performed using ImageJ (Fiji), as described before (*25*).

#### MRI in epilepsy patients

Three epilepsy patients undergoing inpatient monitoring as part of their presurgical evaluation (#1: male, 24 years old | #2: male, 34 years old | #3: female, 34 years old) at the Department of Epileptology at the University of Bonn Medical Center were postictally and interictally scanned in a 3T MAGNETOM Trio (Siemens Healthineers®) applying quantitative MRI. As described (*29*), we acquired quantitative T1 relaxation time maps (qT1) after both ictal and interictal injection of gadolinium-based contrast agent and an additional pre-contrast qT1 scan. The injection-to-MRI time intervals were ensured to be equal for both postictal and interictal scans. Thus, each postictal qT1 volume could be contrasted with interictal qT1 volumes acquired with the same injection-to-MRI time interval. The postictal enhancement of contrast agent was quantified by subtracting postictal qT1 from interictal qT1 and the resulting ΔT1 was used as a surrogate imaging marker of peri-ictal blood-brain barrier dysfunction. Seizure duration, injection latency, and seizure type was determined retrospectively based on the video-EEG-recordings (see legend to Fig. 6C for details). The basic processing steps of the qT1 data were performed using the FMRIB Software Library (FSL) and custom-built scripts written in MATLAB (MathWorks®). The resulting volumes were processed using the standard pipeline of the Freesurfer image analysis suite (http://surfer.nmr.mgh.harvard.edu/). Resulting anatomical labels were extracted and ΔT1 values were averaged across the respective labels. Additional peri-lesional labels were manually defined for patient #2. The average values of labels were color-coded and colored labels were superimposed on the anatomical brain volume.

#### Western blotting

Additional mice were used to assess the extravasation of albumin and IgG into the brain parenchyma (*24*) or the effects of Ant-134 on Limk1 levels (*11*) after status epilepticus. Protein was extracted from perfused brain samples, subjected to SDS-PAGE, transferred to nitrocellulose membranes and incubated with primary antibodies to the following: Albumin (1:1,000), Mouse IgG (1:2,000) (both from Cell Signaling Technology), Limk1 (1:1,000, 3842,from Cell Signaling Technology), β-actin (1:2,000, AC40, Sigma-Aldrich) and α-tubulin (1:1,000, sc-8035, Santa Cruz Biotechnology). Membranes were then incubated with horseradish peroxidase– conjugated secondary antibodies (Jackson ImmunoResearch), and bands were visualized using Supersignal West Pico Chemiluminescence Substrates (Pierce). Images were captured using a Fuji-Film LAS-400, and densitometry was performed using AlphaEaseFC4.0 gel-scanning integrated optical density software (Alpha Innotech). KA+Scr IP and KA+Scr IV were grouped for statistical purposes (KA+Scr). Results are reported as increased protein level (n fold change) in comparison to control samples.

#### Quantitative real-time PCR

Total RNA was isolated from ipsilateral cortex or hippocampus samples using Trizol (Invitrogen), as described previously (*11*). For mRNA qPCR, 1 µg of total RNA was used to generate complementary DNA (cDNA) by reverse transcription using SuperScript III reverse transcriptase enzyme (Invitrogen). Detection of Actin and Limk1 cDNA was performed by SYBR-Green based quantitative Real-time PCR in 20 μL reaction on a Lightcycler 2.0 Carousel based system (Roche™). Reactions were incubated for 15 min at 95°C, followed by 50 cycles of 15 s at 94°C, 20 s at 55°C and 40 s at 72°C (annealing and extension) and then 15 s at 65° (melting). For each sample a Cycle Threshold (Ct) value was determined and then a relative quantification following the ΔΔCt method was carried out using actin as reference gene, all the values obtained was then normalized to the non-transfected samples. The following primer pairs were used: Limk1: fwd 5’-TTATCGGGCGTGTGAATGCA-3’; rev 5’-ACCAGACAAGTGCATGCGAA-3’; House-keeping gene beta-actin: fwd 5’-TAGGCACCAGGGTGATG-3’, rev 5’-GCCATGTTCAATGGGGTACT-3’.

For miRNA, 250 ng of RNA was reverse transcribed using stem-loop specific primers for mmu-miR-134 and real-time quantitative PCR was carried out on a 7900HT Fast Realtime System (Applied Biosystems) using TaqMan miRNA assays. Specific primers were purchased from Sigma and were: RNU19 (Applied Biosystem miRNA assay ID 001186) and miR-134 (Applied Biosystem miRNA assay ID 000395). Expression of RNU19 was used for normalization and relative fold change determined using the ^ΔΔ^CT method (*11*).

#### In situ hybridization

Non-radioactive ISH was performed as described previously (*11*). For detection of Ant-134, sections were prepared from mice killed 24 h or 3 months after status epilepticus. Ant-134-injected mice (*n* = 3/group) were transcardially perfused as before and either the whole brain removed and snap-frozen (24 h samples) or free-floating sections prepared after perfusion fixation (4% PFA). Hybridisation was performed using 5 nM of double-DIG (3’ and 5’) - LNA probe for 134 inhibitor probe (Exiqon) or LNA-DIG Scramble inhibitor (“sense”) probe overnight at 55°C. Slides were washed in SSC buffer, blocked and incubated with anti-DIG antibody overnight followed by 5-bromo-4-chloro-3-indolyl phosphate (BCIP) and nitrobluetetrazolium (NBT) substrates (NBT/BCIP stock solution, Roche Diagnostics), after which staining was stopped by PBS rinses and slides mounted with Vectashield (VectorLabs). Images were acquired with a bright-field microscope and processed on ImageJ, as before (*11*).

Studies using human brain tissue sections were reviewed and approved by the RCSI and Beaumont hospital ethics committees (13/75). Consent was obtained according to the Declaration of Helsinki from all participants. Fresh frozen specimens were from individuals without known neurological disease or patients with intractable temporal lobe epilepsy with or without hippocampal sclerosis (Table S2; n = 3/group) and processed as described, with modifications (*11*). Briefly, 16 μm thick sections were fixed, acetylated and treated with proteinase K. Hybridisation was performed with 10 nM of double-DIG (3’ and 5’) - LNA probe for human-miR-134 (Exiqon) or LNA-DIG Scramble probe overnight at 50^º^C. Slides were washed in SSC buffer and blocked with 10% foetal calf serum, then incubated with anti-digoxigenin-AP Fab fragments (1:2,500, Roche Diagnostics). Slides were treated with NBT/BCIP and staining was stopped by washing in PBS. Slides were mounted using vectashield (VectorLabs) and images acquired with brightfield microscope and processed on ImageJ.

For FISH, after blocking, slides were co-incubated with anti-Digoxigenin-POD (1;500, Roche Diagnostics) and NeuN (1;400, Millipore) or GFAP (1;1000, Dako Cytomation) antibody. Signal was amplified using TSA™ Cyanine 3 System (1;50 in amplification diluent, PerkinElmer). After rinsing, slides were incubated with secondary antibody (Alexafluor 488, Invitrogen). Nuclei were stained with DAPI and slides were mounted using ProLong Gold (Life Technologies). Images were acquired using a confocal microscope (LSM880, Zeiss).

#### Immunohistochemistry

Immunohistochemistry was performed on fresh frozen coronal sections obtained after long-term EEG experiments. Slices were air-dried and post-fixed, rinsed and permeabilised in 3% Triton-X-100/PBS. Next, sections were blocked in 5% goat serum followed by overnight incubation with primary antibody (1:400; NeuN, GFAP, Iba-1 and Parvalbumin (all from Cell Signalling Techn.) Slides were washed again and incubated with fluorescently-labelled secondary antibodies, AlexaFluor 488 and AlexaFluor 546 (1:400) and Hoechst was added. Staining was imaged using a Hamamatsu Orca 285 camera attached to a Nikon 2000s epifluorescence microscope (Micron Optical). Positive cells were quantified per hippocampal subfield, and were the average of two adjacent sections by an observer blinded to the experimental treatment. Neuronal damage (NeuN) was assessed based on a graded five-point system under 40× magnification: level 0, no damage; level 1, 0–25%; level 2, 25–50%; level 3, 50–75%; level 5, 75% cell loss.

#### Transcriptomics analysis (microarray)

A separate set of mice underwent surgical and EEG procedures for acute assessments and status epilepticus followed by Ant-134 (30mg/kg.^-1^, IP) or Scr control, as above. Twenty-four hours later, mice were transcardially perfused with ice-cold PBS and the ipsilateral hippocampus was snap frozen. Total RNA was isolated as previously described and miR-134 mRNA targets were analysed by microarray. Microarrays used were Mouse Whole Genome GE 4×44K v1 (Agilent Technologies, Belgium) representing 41174 *M. musculus* 60-mer probes in a 4×44K layout. Microarray experiments were performed 8 times: - for each sample group, two samples were labelled with cy5 and co-hybridized with reference pool RNA labelled with cy3, and two samples were labelled and hybridized in the opposite way. RNA amplifications and labelling were performed on an automated system (Caliper Life Sciences NV/SA, Belgium) with 3 μg total RNA from each sample. Hybridizations were done on a HS4800PRO system with QuadChambers (Tecan Benelux B.V.B.A.) using 1000 ng labelled cRNA per channel according to van Wageningen et al (*39*). Hybridized slides were scanned on an Agilent scanner (G2565BA) at 100% laser power, 30% PMT. After automated data extraction using Imagene 8.0 (BioDiscovery), Loess normalization was performed on mean spot-intensities and gene-specific dye bias was corrected based on a within-set estimate. Differential expression p-values (Benjamini-Hochberg FDR correction) were calculated using the limma package in R V2.12. For mRNA with multiple isoforms, we selected the maximal p-value and associated fold change. Two mRNA (Hba-b2, Hbb-b2) with log2FC < −1.5 were removed from the plotted results for clarity. Data will be deposited to GEO on acceptance.

#### Structural modelling of Ant-134

The 3D model of Ant-134 and Ant-134 bound to miR-134 were generated using Molecular Operating Environment, CCG Montreal.

#### Statistical analysis

Statistical analysis was performed using GraphPad Prism and Stata Release 15 software, and a probability of P < 0.05 was considered significant. Two-Way ANOVA was used to analyse the relative expression of albumin and mouse IgG. One-Way ANOVA and Bonferroni ‘s multiple comparisons test was used to ratify the successful delivery of Ant-134 by analysing the relative levels of miR-134 and Limk1 expression. Negative binomial regression was used to compare the daily incidence of seizures (SRS) between the Ant-134-treated and control groups. Ordinal logistic regression was used to examine the relationship of time (days) on the number of recorded seizures. An interaction term was used to determine if the relationship differed between the scramble and Ant-134 groups. Differences between Ant-134 and Scr-injected animals in immunohistochemical cell counts were analysed by Student’s t-test. Data are expressed as mean + S.E.M. In all cases, robust variance estimation was used to adjust standard errors for clustering of observations within animals.

## Acknowledgements

The authors would like to thank Eva Jimenez-Mateos and Tobias Engel for advice on technical aspects. This publication has emanated from research supported in part by a research grant from Science Foundation Ireland (SFI) under Grant Number 16/RC/3948 and co-funded under the European Regional Development Fund and by FutureNeuro industry partners. The authors also acknowledge funding from the Health Research Board Ireland (HRA-POR-2013-325, to D.C.H.) and Science Foundation Ireland (13/IA/1891, 11/TIDA/B1988, to D.C.H.), fellowships from the Brazilian National Council for Scientific and Technological Development (CNPq; to L.F.A.S.), the Dutch Epilepsiefonds (WAR 12-08, 15-05) (to R.J.P.), the Royal Society (to S.S.), the Irish Research Council and Science Foundation Ireland (17/TIDA/5002, to C.R.R.), CURE (D.C.H.) and the European Union Seventh Framework Programme (FP7/2007-2013) under grant agreement n° 602130. We gratefully acknowledge the NIH NeuroBioBank and University of Maryland Brain and Tissue Bank for autopsy material. The role of the Brain and Tissue Bank is to distribute tissue, and therefore, cannot endorse the studies performed or the interpretation of results.

## Author contributions

CRR performed *in vivo* acute and long-term antagomir studies and data analysis in mice. LFAS performed immunohistochemistry and overall data analysis. VV and RJP performed *in situ* hybridization and microarray assays. MR, CRR and ASR performed in vitro characterization of GapmeRs. BD and TR performed and analysed human MRI. BLC and CRR performed two-photon microscopy. NMCC, CM and JP performed bioinformatic analysis. GPB and ASR performed qPCRs and western blotting for miR-134 and Limk1 and data analysis. AB and CRR performed Evans Blue analysis, Albumin and mouse IgG western blotting, and FITC histology experiments. CG and MC performed the MRI imaging in mice. MB performed the Ant-134 structure modelling and predictions. RMC assisted with experimental design and statistical analysis. CRR and DCH conceived and designed the animal studies and DCH wrote the initial manuscript. All authors contributed and approved the final manuscript. Authors state that all data associated with this study are available in the main text or the supplementary materials.

## References

1. A. A. Levin, Treating Disease at the RNA Level with Oligonucleotides. N Engl J Med 380, 57–70 (2019).

2. O. Khorkova, C. Wahlestedt, Oligonucleotide therapies for disorders of the nervous system. Nat Biotechnol 35, 249–263 (2017).

3. B. Obermeier, R. Daneman, R. M. Ransohoff, Development, maintenance and disruption of the blood-brain barrier. Nat Med 19, 1584–1596 (2013).

4. O. Devinsky, A. Vezzani, T. J. O’Brien, N. Jette, I. E. Scheffer, M. de Curtis, P. Perucca, Epilepsy. Nat Rev Dis Primers 4, 18024 (2018).

5. W. Loscher, The holy grail of epilepsy prevention: Preclinical approaches to antiepileptogenic treatments. Neuropharmacology, (2019).

6. D. C. Henshall, H. M. Hamer, R. J. Pasterkamp, D. B. Goldstein, J. Kjems, J. H. Prehn, S. Schorge, K. Lamottke, F. Rosenow, MicroRNAs in epilepsy: pathophysiology and clinical utility. Lancet Neurol 15, 1368–1376 (2016).

7. E. McNeill, D. Van Vactor, MicroRNAs Shape the Neuronal Landscape. Neuron 75, 363–379 (2012).

8. R. C. McKiernan, E. M. Jimenez-Mateos, I. Bray, T. Engel, G. P. Brennan, T. Sano, Z. Michalak, C. Moran, N. Delanty, M. Farrell, D. O’Brien, R. Meller, R. P. Simon, R. L. Stallings, D. C. Henshall, Reduced mature microRNA levels in association with Dicer loss in human temporal lobe epilepsy with hippocampal sclerosis. PLoS One 7, e35921 (2012).

9. S. S. Hebert, A. S. Papadopoulou, P. Smith, M. C. Galas, E. Planel, A. N. Silahtaroglu, N. Sergeant, L. Buee, B. De Strooper, Genetic ablation of Dicer in adult forebrain neurons results in abnormal tau hyperphosphorylation and neurodegeneration. Hum Mol Genet 19, 3959–3969 (2010).

10. C. L. Tan, J. L. Plotkin, M. T. Veno, M. von Schimmelmann, P. Feinberg, S. Mann, A. Handler, J. Kjems, D. J. Surmeier, D. O’Carroll, P. Greengard, A. Schaefer, MicroRNA-128 governs neuronal excitability and motor behavior in mice. Science 342, 1254–1258 (2013).

11. E. M. Jimenez-Mateos, T. Engel, P. Merino-Serrais, R. C. McKiernan, K. Tanaka, G. Mouri, T. Sano, C. O’Tuathaigh, J. L. Waddington, S. Prenter, N. Delanty, M. A. Farrell, D. F. O’Brien, R. M. Conroy, R. L. Stallings, J. Defelipe, D. C. Henshall, Silencing microRNA-134 produces neuroprotective and prolonged seizure-suppressive effects. Nat Med 18, 1087–1094 (2012).

12. C. R. Reschke, L. Fernando, B. A. Norwood, K. Senthilkumar, G. Morris, A. Sanz-Rodriguez, R. Conroy, L. Costard, V. Neubert, S. Bauer, S. Schorge, R. J. Pasterkamp, F. Rosenow, D. C. Henshall, Potent anti-seizure effects of locked nucleic acid antagomirs targeting miR-134 in multiple mouse and rat models of epilepsy. Mol Thera 6, 45–56 (2017).

13. X. Gao, M. Guo, D. Meng, F. Sun, L. Guan, Y. Cui, Y. Zhao, X. Wang, X. Gu, J. Sun, S. Qi, Silencing MicroRNA-134 Alleviates Hippocampal Damage and Occurrence of Spontaneous Seizures After Intraventricular Kainic Acid-Induced Status Epilepticus in Rats. Front Cell Neurosci 13, 145 (2019).

14. G. M. Schratt, F. Tuebing, E. A. Nigh, C. G. Kane, M. E. Sabatini, M. Kiebler, M. E. Greenberg, A brain-specific microRNA regulates dendritic spine development. Nature 439, 283–289 (2006).

15. J. Gao, W. Y. Wang, Y. W. Mao, J. Graff, J. S. Guan, L. Pan, G. Mak, D. Kim, S. C. Su, L. H. Tsai, A novel pathway regulates memory and plasticity via SIRT1 and miR-134. Nature 466, 1105–1109 (2010).

16. P. Gaughwin, M. Ciesla, H. Yang, B. Lim, P. Brundin, Stage-specific modulation of cortical neuronal development by Mmu-miR-134. Cereb Cortex 21, 1857–1869 (2011).

17. J. Elmen, M. Lindow, S. Schutz, M. Lawrence, A. Petri, S. Obad, M. Lindholm, M. Hedtjarn, H. F. Hansen, U. Berger, S. Gullans, P. Kearney, P. Sarnow, E. M. Straarup, S. Kauppinen, LNA-mediated microRNA silencing in non-human primates. Nature 452, 896–899 (2008).

18. E. M. Jimenez-Mateos, T. Engel, P. Merino-Serrais, I. Fernaud-Espinosa, N. Rodriguez-Alvarez, J. Reynolds, C. R. Reschke, R. M. Conroy, R. C. McKiernan, J. deFelipe, D. C. Henshall, Antagomirs targeting microRNA-134 increase hippocampal pyramidal neuron spine volume in vivo and protect against pilocarpine-induced status epilepticus. Brain Struct Funct 220, 2387–2399 (2015).

19. E. M. Straarup, N. Fisker, M. Hedtjarn, M. W. Lindholm, C. Rosenbohm, V. Aarup, H. F. Hansen, H. Orum, J. B. Hansen, T. Koch, Short locked nucleic acid antisense oligonucleotides potently reduce apolipoprotein B mRNA and serum cholesterol in mice and non-human primates. Nucleic Acids Res 38, 7100–7111 (2010).

20. E. A. van Vliet, S. da Costa Araujo, S. Redeker, R. van Schaik, E. Aronica, J. A. Gorter, Blood-brain barrier leakage may lead to progression of temporal lobe epilepsy. Brain 130, 521–534 (2007).

21. D. Shlosberg, M. Benifla, D. Kaufer, A. Friedman, Blood-brain barrier breakdown as a therapeutic target in traumatic brain injury. Nat Rev Neurol 6, 393–403 (2010).

22. S. Lublinsky, S. Major, V. Kola, V. Horst, E. Santos, J. Platz, O. Sakowitz, M. Scheel, C. Dohmen, R. Graf, H. Vatter, S. Wolf, P. Vajkoczy, I. Shelef, J. Woitzik, P. Martus, J. P. Dreier, A. Friedman, Early blood-brain barrier dysfunction predicts neurological outcome following aneurysmal subarachnoid hemorrhage. EBioMedicine 43, 460–472 (2019).

23. E. M. Jimenez-Mateos, M. Arribas-Blazquez, A. Sanz-Rodriguez, C. Concannon, L. A. Olivos-Ore, C. R. Reshcke, C. M. Mooney, C. Mooney, E. Lugara, J. Morgan, E. Langa, A. Jimenez-Pacheco, L.-F. Almeida-Silva, G. B. Mesuret, D., M. T. Miras-Portugal, M. Letavic, A. R. Artalejo, A. Bhattacharya, M. Diaz-Hernandez, D. C. Henshall, T. Engel, MicroRNA targeting of the P2X7 purinoceptor opposes a contralateral epileptogenic focus in the hippocampus. Scientific Reports 5, e17486 (2015).

24. Z. Michalak, T. Sano, T. Engel, S. Miller-Delaney, M. Lerner-Natoli, D. C. Henshall, Spatio-temporally restricted blood-brain barrier disruption after intra-amygdala kainic acid-induced status epilepticus in mice. Epilepsy Res 103, 167–179 (2013).

25. J. Keaney, D. M. Walsh, T. O’Malley, N. Hudson, D. E. Crosbie, T. Loftus, F. Sheehan, J. McDaid, M. M. Humphries, J. J. Callanan, F. M. Brett, M. A. Farrell, P. Humphries, M. Campbell, Autoregulated paracellular clearance of amyloid-beta across the blood-brain barrier. Sci Adv 1, e1500472 (2015).

26. T. Li, G. Ren, T. Lusardi, A. Wilz, J. Q. Lan, T. Iwasato, S. Itohara, R. P. Simon, D. Boison, Adenosine kinase is a target for the prediction and prevention of epileptogenesis in mice. J Clin Invest 118, 571–582 (2008).

27. X. Jiang, M. Lachance, E. Rossignol, Involvement of cortical fast-spiking parvalbumin-positive basket cells in epilepsy. Prog Brain Res 226, 81–126 (2016).

28. R. J. Buono, F. W. Lohoff, T. Sander, M. R. Sperling, M. J. O’Connor, D. J. Dlugos, S. G. Ryan, G. T. Golden, H. Zhao, T. M. Scattergood, W. H. Berrettini, T. N. Ferraro, Association between variation in the human KCNJ10 potassium ion channel gene and seizure susceptibility. Epilepsy Res 58, 175–183 (2004).

29. T. Ruber, B. David, G. Luchters, R. D. Nass, A. Friedman, R. Surges, T. Stocker, B. Weber, R. Deichmann, G. Schlaug, E. Hattingen, C. E. Elger, Evidence for peri-ictal blood-brain barrier dysfunction in patients with epilepsy. Brain 141, 2952–2965 (2018).

30. C. Gross, X. Yao, T. Engel, L. Xing, S. W. Danielson, K. T. Thomas, E. M. Jimenez-Mateos, D. C. Henshall, G. J. Bassell, MicroRNA-mediated downregulation of the potassium channel Kv4.2 contributes to seizure onset. Cell Rep 17, 37–45 (2016).

31. G. Morris, G. P. Brennan, C. R. Reschke, D. C. Henshall, S. Schorge, Spared CA1 pyramidal neuron function and hippocampal performance following antisense knockdown of microRNA-134. Epilepsia in press, (2018).

32. S. Sambandan, G. Akbalik, L. Kochen, J. Rinne, J. Kahlstatt, C. Glock, G. Tushev, B. Alvarez-Castelao, A. Heckel, E. M. Schuman, Activity-dependent spatially localized miRNA maturation in neuronal dendrites. Science 355, 634–637 (2017).

33. J. Wetherington, G. Serrano, R. Dingledine, Astrocytes in the epileptic brain. Neuron 58, 168–178 (2008).

34. H. L. Janssen, H. W. Reesink, E. J. Lawitz, S. Zeuzem, M. Rodriguez-Torres, K. Patel, A. J. van der Meer, A. K. Patick, A. Chen, Y. Zhou, R. Persson, B. D. King, S. Kauppinen, A. A. Levin, M. R. Hodges, Treatment of HCV infection by targeting microRNA. N Engl J Med 368, 1685–1694 (2013).

35. X. Wang, Y. Luo, S. Liu, L. Tan, S. Wang, R. Man, MicroRNA-134 plasma levels before and after treatment with valproic acid for epilepsy patients. Oncotarget 8, 72748–72754 (2017).

36. H. McArdle, E. M. Jimenez-Mateos, R. Raoof, E. Carthy, D. Boyle, H. ElNaggar, N. Delanty, H. Hamer, M. Dogan, T. Huchtemann, P. Krtvelyessy, F. Rosenow, R. J. Forster, D. C. Henshall, E. Spain, “TORNADO” - Theranostic One-Step RNA Detector; microfluidic disc for the direct detection of microRNA-134 in plasma and cerebrospinal fluid. Sci Rep 7, 1750 (2017).

37. D. M. Kullmann, S. Schorge, M. C. Walker, R. C. Wykes, Gene therapy in epilepsy-is it time for clinical trials? Nat Rev Neurol 10, 300–304 (2014).

38. J. Schindelin, I. Arganda-Carreras, E. Frise, V. Kaynig, M. Longair, T. Pietzsch, S. Preibisch, C. Rueden, S. Saalfeld, B. Schmid, J. Y. Tinevez, D. J. White, V. Hartenstein, K. Eliceiri, P. Tomancak, A. Cardona, Fiji: an open-source platform for biological-image analysis. Nat Methods 9, 676–682 (2012).

39. S. van Wageningen, P. Kemmeren, P. Lijnzaad, T. Margaritis, J. J. Benschop, I. J. de Castro, D. van Leenen, M. J. Groot Koerkamp, C. W. Ko, A. J. Miles, N. Brabers, M. O. Brok, T. L. Lenstra, D. Fiedler, L. Fokkens, R. Aldecoa, E. Apweiler, V. Taliadouros, K. Sameith, L. A. van de Pasch, S. R. van Hooff, L. V. Bakker, N. J. Krogan, B. Snel, F. C. Holstege, Functional overlap and regulatory links shape genetic interactions between signaling pathways. Cell 143, 991–1004 (2010).

40. J. J. Engel, P. C. Van Ness, T. B. Rasmussen, in Surgical treatment of the epilepsies, Engel J, Ed. (Raven Press, New York, 1993), pp. 609–621.

