## Supplementary file for "Potent and lasting seizure suppression by systemic delivery of antagomirs targeting miR-134 timed with blood-brain barrier disruption"

**Figure S1**

Blood brain barrier disruption after intra-amygdala kainic acid-induced status epilepticus.

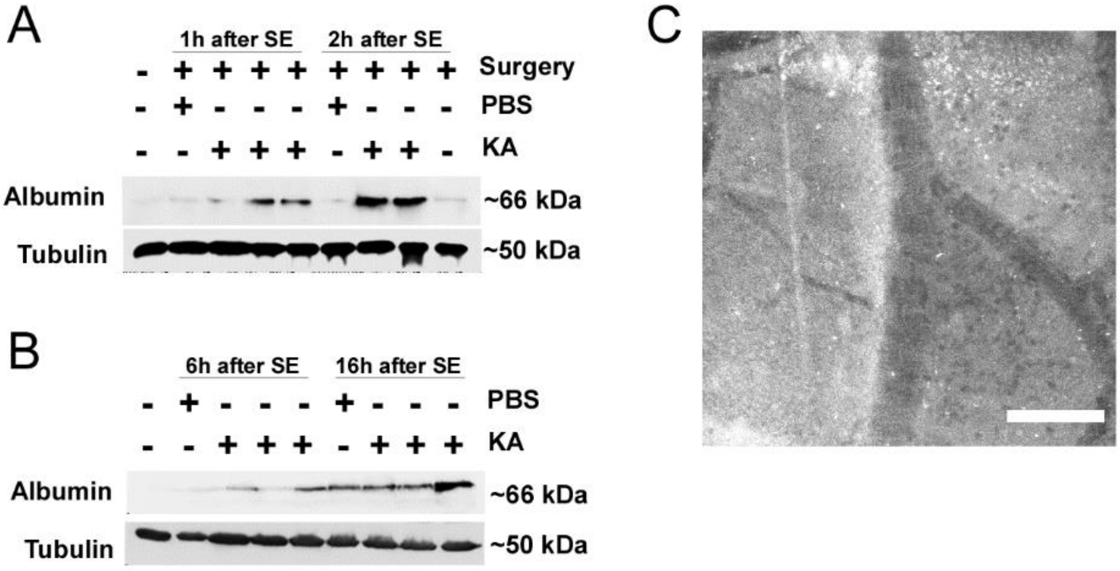


**Figure S1.** (A-B) Representative western blots showing the extravasation of Albumin into the brain mouse hippocampus at 1h, 2h, 6h and 16h after status epilepticus. (C) Extravasation of FITC-dextran into the brain parenchyma after KA-induced status epilepticus. Maximum image projection images of multiphoton Z stack Images at 2 hours after the first, and 1 hour after a second FITC-dextran injection. Scale bar represents 100µm.

### Figure S2

Additional quantification of SRS during long-term EEG monitoring in Ant-134 and Scr mice

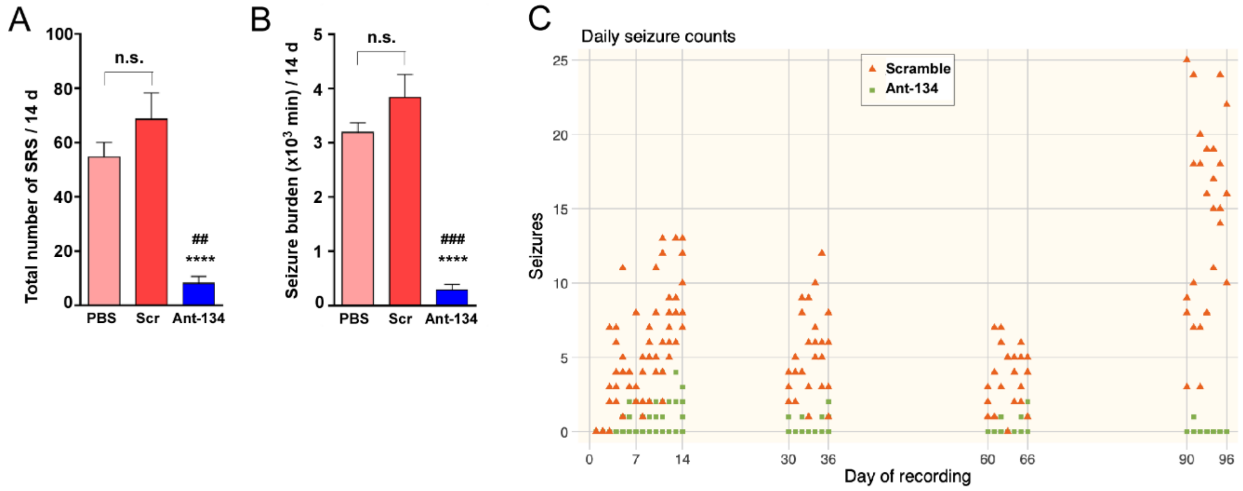


**Figure S2.** (A-B) Graphs show no differences between either controls PBS or Scramble (Scr) in the total number of (A) spontaneous recurrent seizures (SRS) or (B) seizure burden (total time spent in ictal activity in minutes). On the other hand, Ant-134-injected mice presented a significant reduction in both measure parameters in comparison with either controls. ## P<0.01, ### P<0.001 when compared with PBS. **** P<0.0001 when compared with Scramble by One-Way ANOVA. (C) SRS distribution over all activation periods, plotted individually per day per treatment group.

**Figure S3** Additional histological findings in Ant-134-treated mice.

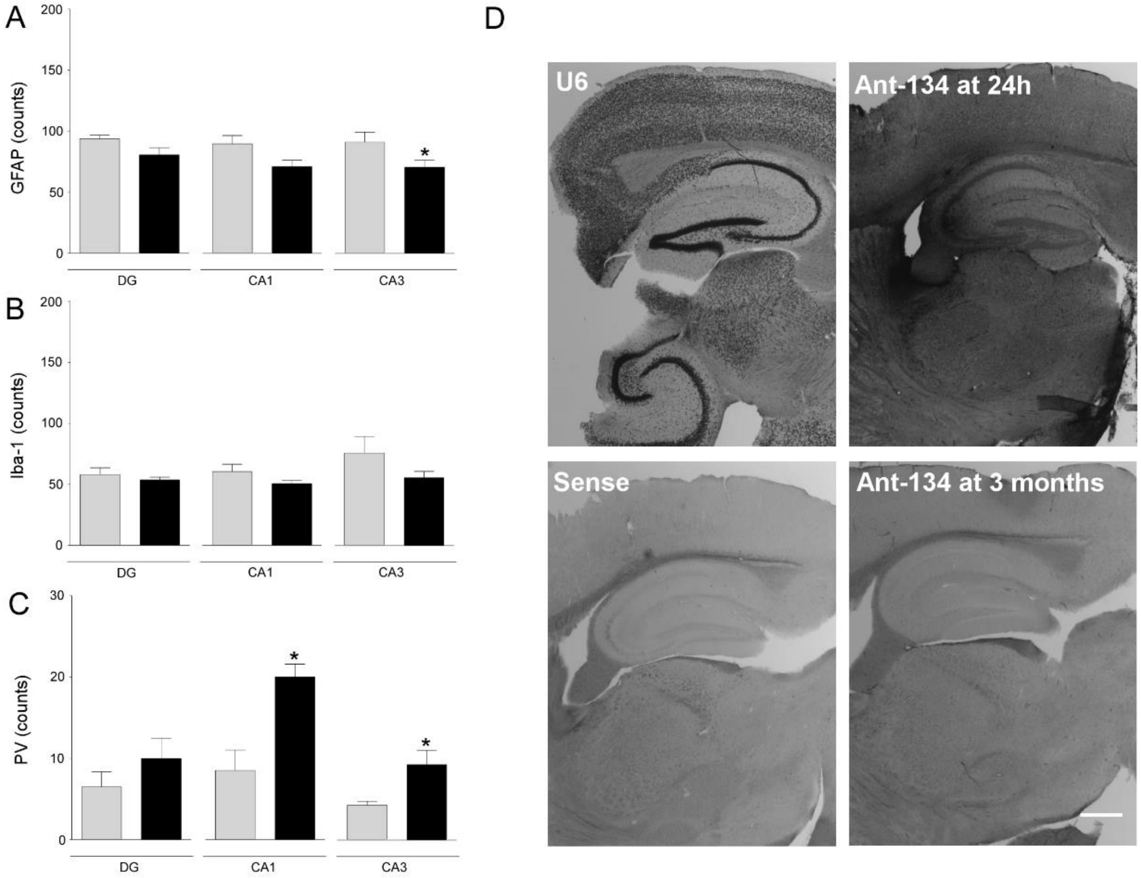

**Figure S3.** (A-C) Counts of GFAP (A), Iba-1 (B) and Parvalbumin (C) positive cells per hippocampal subfield (DG, CA1 and CA3). Grey and black bars represent Scramble and Ant-134 groups, respectively, as Mean ± SEM. * $P < 0.05$ when compared with Scramble by One-Way ANOVA. (D) *In situ* hybridization shows the presence of Ant-134 in the mouse brain at 24h after its IP injection [(30 $mg.kg^{-1}$), picture top right], which was no longer present in the brain at 3 months time point [(end of EEG recording “Period 4”) picture bottom right]. Sense [(scramble inhibitor probe), picture bottom left] and U6 (picture top left) were used as negative and positive controls, respectively. Scale bar represents 500 µm.

**Figure S4** LIMK-GapmeR *in vitro* characterization.

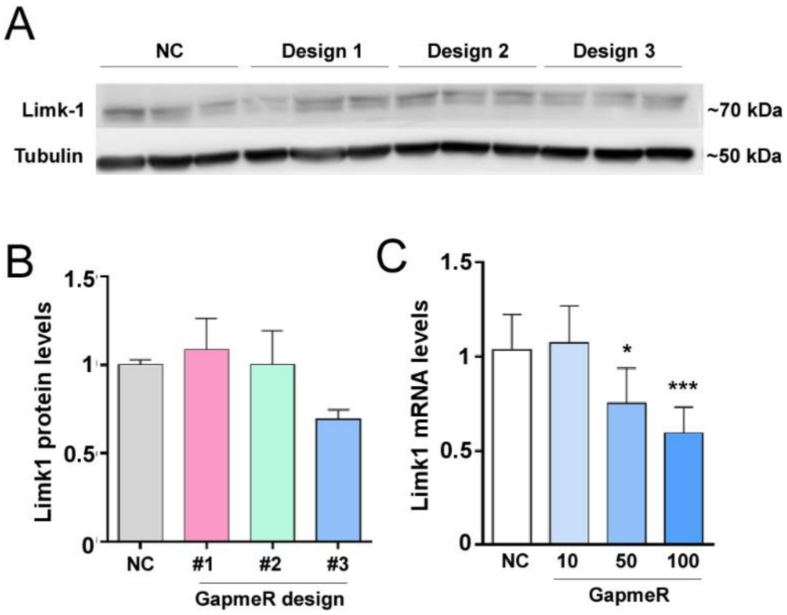


**Figure S4.** (A-C) Three LIMK-GapmeR were *in silico* predicted and designed for transfection in N2A cells (240,000 cells/well; 100 nM). Protein was extracted after 24h for Western Blot analysis of Limk1 expression, corrected to α-tubulin. (A) Western blot depicts the expression of Limk1 and α-tubulin proteins for LIMK-GapmeR Design 1 (#1), Design 2 (#2) and Design 3 (#3). (B) Graph shows Limk1 relative protein levels per GapmeR predicted design in comparison with negative control [(NC); non-transfected sample]. (C) Graph shows Limk1 relative protein expression after transfection with different doses of a Premium (HPLC purified) version of LIMK-GapmeR design 3 (#3). *P<0.05 and ***P<0.001 when compared to NC by One-Way ANOVA.

**Figure S5** Cellular distribution of miR-134 in pharmacoresistant epilepsy patients.

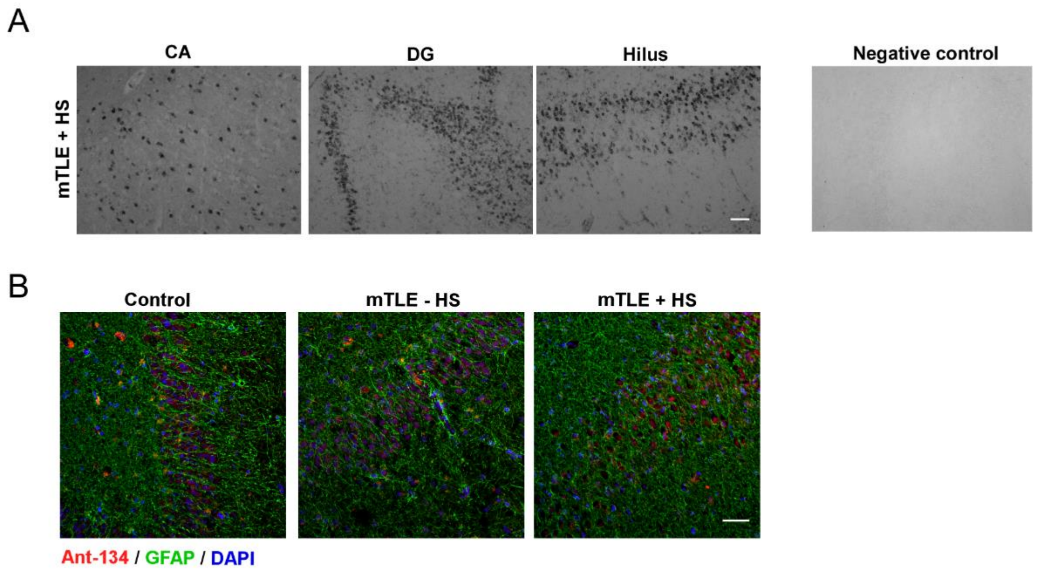


**Figure S5.** (A) Non-radioactive *in situ* hybridization (ISH) shows the expression of miR-134 in the CA, DG and Hilus of TLE patients with hippocampal sclerosis (mTLE + HS). Control images (non-epilepsy patients) are shown in Figure 6 of the main manuscript, and negative control is the scramble probe. Scale bar represents 100µm. (n=3-4/group). (B) Representative pictures show that miR-134 does not co-localizes with astrocytes (GFAP) in brain tissue samples from mTLE ± HS in comparison with non-epilepsy controls (DG; fluorescence ISH; scale bar represents 50µm; n=3-4/group).

**Table S1** Number of spontaneous recurrent seizures (SRS) and seizure-free days per treatment group per EEG period.

| | | Scramble | Ant-134 |
|---|---|---|---|
| **Period 1** | SRS number* | 68.6 | 8.16 |
| | Seizure-free days* | 0.4 | 7 |
| **Period 2** | SRS number ** | 35.5 | 1.33 |
| | Seizure-free days** | - | 5.83 |
| **Period 3** | SRS number ** | 25.75 | 1.4 |
| | Seizure-free days** | - | 5.8 |
| **Period 4** | SRS number ** | 100 | 0.2 |
| | Seizure-free days** | - | 6.8 |

*Average per group for 12 days of recording (from the 3rd to 14th day) of activation period 1. Mice from either groups did not present spontaneous recurrent seizures (SRS) before day 3.

**Average per group for 7 days of recording per each activation period.

### Table S2

Clinical details from control and mTLE patients used for *in situ* hybridization experiments.

| **Sample** | **Age** | **Sex** | **PMD** | **Age of onset** | **Years of epilepsy** | **AED's** |
|---|---|---|---|---|---|---|
| Control | 71 | M | 9h | NA | NA | NA |
| Control | 94 | F | 4h | NA | NA | NA |
| Control | 58 | M | 7h | NA | NA | NA |
| mTLE-HS | 46 | M | NA | 33 | 13 | CBZ, VPA, TPR |
| mTLE-HS | 40 | F | NA | 17 | 23 | LEV, LTG, CBZ |
| mTLE-HS | 45 | M | NA | 18 | 27 | LTG, PHT |
| mTLE-HS | 50 | F | NA | 36 | 14 | LTG,CBZ |
| mTLE+HS | 43 | M | NA | 13 | 30 | OXC, CLO,LEV |
| mTLE+HS | 44 | F | NA | 8 | 36 | GBP, CLO |
| mTLE+HS | 12 | F | NA | 5.5 | 6.5 | CLO, LEV, LTG, CBZ |
| mTLE+HS | 15 | M | NA | 12 | 3 | DZP,LTG, OXC |

**Table S2.** Clinical data available and details of anti-epileptic drug (AED) medication used for TLE patients. LTG lamotrigine, PHT phenytoin, CBZ carbamazepine, GBP gabapentin, LEV levetiracetam, OXC oxcarbazepine, CLO clobazam, DZP diazepam, LZP lorazepam, SER Seroquel, PGB pregabaline, RES restoril.
